# Alterations in neuronal physiology, development, and function associated with a common duplication of chromosome 15 involving *CHRNA7*

**DOI:** 10.1101/2020.01.28.922187

**Authors:** Kesavan Meganathan, Ramachandran Prakasam, Dustin Baldridge, Paul Gontarz, Bo Zhang, Fumihiko Urano, Azad Bonni, James E. Huettner, John N. Constantino, Kristen L. Kroll

## Abstract

**Background:** Copy number variants at chromosome 15q13.3 contribute to liability for multiple intellectual and developmental disabilities including Autism Spectrum Disorder (ASD). Individuals with duplications of this interval, which includes the gene *CHRNA7*, have multiple psychiatric disorders with widely variable penetrance. However, the basis of such differential affectation remains uncharacterized.

**Methods:** Induced pluripotent stem cell (iPSC) models were generated from two first degree relatives with the same 15q13.3 duplication, a boy with distinct features of autism and emotional dysregulation (the affected proband, AP) and his clinically unaffected mother (the UM). These models were compared to unrelated control subjects lacking this duplication (UC, male and female). iPSC-derived neural progenitors and cortical neuroids consisting of cortical excitatory and inhibitory neurons were used to model potential contributors to neuropsychiatric impairment.

**Results:** The AP-derived model uniquely exhibited disruptions of cellular physiology and neurodevelopment not observed in either the UM or the unrelated male and female controls. These included enhanced neural progenitor proliferation but impaired neuronal differentiation, maturation, and migration, and increased endoplasmic reticulum (ER) stress. Both the AP model’s neuronal migration deficit and elevated ER stress could be selectively rescued by different pharmacologic agents. Neuronal gene expression was also specifically dysregulated in the AP, including reduced expression of genes related to behavior, psychological disorders, neuritogenesis, neuronal migration, and WNT, axonal guidance, and GABA receptor signaling. Interestingly, the UM model exhibited upregulated expression of genes in many of these same pathways, by comparison with both the AP and UC models, suggesting that cell intrinsic molecular compensation could have contributed to the lack of neurodevelopmental phenotypes in the UM model. However, by contrast with the AP-specific neurodevelopmental phenotypes, both the AP- and UM-derived neurons exhibited shared alterations of neuronal function, including increased action potential firing and elevated cholinergic activity, consistent with increased homomeric CHRNA7 channel activity.

**Conclusion:** Together, these data define both affectation-specific phenotypes seen only in the AP, as well as abnormalities observed in both individuals with *CHRNA7* duplication, the AP and UM, but not in UC-derived neurons. This is, to our knowledge, the first study to use a human stem cell-based platform to study the basis of variable affectation in cases of 15q13.3 duplication at the cellular, molecular, and functional levels. This work suggests potential approaches for suppressing abnormal neurodevelopment or physiology that may contribute to severity of affectation. Some of these AP-specific neurodevelopmental anomalies, or the functional anomalies observed in both 15q13.3 duplication carriers (the AP and UM), could also contribute to the variable phenotypic penetrance seen in other individuals with 15q13.3 duplication.

## Introduction

Reciprocal copy number variants (CNVs) related to neurodevelopmental and neuropsychiatric disorders including autism spectrum disorder (ASD), schizophrenia, epilepsy, and intellectual disability (ID) result from single copy deletion or duplication of susceptible genomic intervals (1–3). While CNVs involving deletion often cause severe, highly penetrant patient phenotypes, CNVs involving genomic microduplication frequently exhibit much more variable and less penetrant phenotypic expressivity among affected individuals in the population (4–6). Among these, either chromosome 15q13.3 deletion or duplication can result in a range of clinical phenotypes including ASD, ID, mood disorder, language delay, or schizophrenia (7–9). Microdeletions in the 15q13.3 region often result in severe cognitive deficits, behavioral abnormalities, and highly penetrant ASD (7, 10). By comparison, individuals with microduplications at 15q13.3 often exhibit milder phenotypes, including borderline Intellectual Disability (ID), ASD, and attention deficit hyperactivity disorder (ADHD) (7, 11). Notably, sequence gains at 15q13.3 are present in 1.25% of reported ADHD probands, but also in 0.61% of control subjects, suggesting that some individuals tolerate 15q13.3 duplication without clinical affectation, due to its poor phenotypic penetrance (7, 8, 11).

Duplications in the 15q13.3 region of the genome either duplicate only the α-7 nicotinic acetylcholine receptor subunit gene (*CHRNA7*), with or without the first exon of the *OTUD7A* gene, or instead duplicate several genes (*CHRNA7, OTUD7A*, and *UBE3A*) (7, 12–15). *CHRNA7* is a prototypical genetic contributor to complex psychiatric disease: receptors in this protein family form ligand-gated ion channels comprised of five homomeric or heteromeric acetylcholine receptor subunits and are stimulated endogenously by choline and acetylcholine, resulting in flux of the cations calcium, sodium, and potassium (16). Clinical phenotypes associated with *CHRNA7* gene duplication include motor delays, hypotonia, ASD, intellectual disability, schizophrenia, and epilepsy, with the particular phenotypes exhibited varying by individual (7, 15–17). Although the clinical significance of an increased dosage of the CHRNA7 receptor is unclear, no other detectable copy number variation is detected in most reported cases, while clinical data suggests that *CHRNA7* duplication is pathogenic, but with reduced and highly variable penetrance (7, 8, 16).

Human cellular models generated from patient-derived induced pluripotent stem cells (iPSCs) provide a powerful approach for modeling neurodevelopmental disorders, including ASD. In recent years, we and others have characterized cellular phenotypes linked to neurodevelopmental disorders, either in patient iPSC-derived neurons or after introduction of a pathogenic mutation into control iPSCs using CRISPR/Cas9-based gene editing. iPSC-based studies have been conducted to characterize syndromic forms of ASD, de novo cases, and instances where either monogenic or polygenic contributors to disease are indicated (18–22). These studies identified potential contributors to disease, including alterations in gene expression, differential regulation of developmental signaling pathways, and impairment of neurogenesis and synaptogenesis (19–21, 23–26). In addition to identifying phenotypes linked to affectation, some of these studies found specific, disorder-linked targets that were amenable to pharmacological rescue (18, 27, 28).

While CNVs at human 15q13.3 involving *CHRNA7* duplication contribute to a range of neuropsychiatric disorders, the consequences of these CNVs have not been extensively modeled. *CHRNA7* knockout in the mouse, mimicking some 15q13.3 deletions, did not result in behavioral phenotypes (10, 11, 29). *CHRNA7* duplication has not been modeled *in vivo* and the size of 15q13.3 duplications precludes modeling them by genome engineering in rodents. CHRNA7 overexpression in a mouse neuroblastoma cell line altered receptor sensitivity to choline and varenicline (30). Two studies have characterized human iPSC-derived models involving duplications in the chromosome 15q region (28, 31). One study focused on a large duplication of 15q11-q13.1 involving 33 genes; a maternally expressed gene (*UBE3A*) was found to be upregulated in response to duplication of this interval, and this effect could be pharmacologically rescued (28). A second study utilized three iPSC-derived excitatory neuron progenitor cell (NPC) models each from subjects with either 15q13.3 duplication or deletion (31). These models exhibited increased or decreased CHRNA7 mRNA levels, respectively, while both duplication and deletion reduced CHRNA7-dependent calcium flux, indicating diminished channel activity. In the duplication models, this finding was linked to elevated expression of several chaperone genes that control CHRNA7 protein folding and trafficking through the ER to the cell surface; this finding suggested that inefficient chaperoning and accumulation of excess CHRNA7 in the ER, rather than at the cell membrane, could be responsible for the diminished CHRNA7 channel activity observed (31).

Duplications involving CHRNA7 are common contributors to psychopathophysiology, exhibit widely variable phenotypic penetrance, and are difficult to model in animals. While a prior study provided evidence for altered physiology of NPCs carrying *CHRNA7* duplications (31), how this CNV more broadly affects neurodevelopment, global gene expression, and electrophysiological function of neurons had not been evaluated. Potential contributors to the variable phenotypic penetrance exhibited by 15q13.3 duplication carriers were also entirely unknown. Therefore, here we generated iPSC models from two first degree relatives with the same 15q13.3 copy number variant, which duplicates only the *CHRNA7* gene. These subjects include a boy with distinct features of autism and emotional dysregulation (the affected proband, AP) and his clinically unaffected mother (the UM). These models were compared to unrelated male and female control subjects lacking this duplication (the UC-M/UC-F). We used these models to assess the consequences of *CHRNA7* duplication on the development and function of both cortical excitatory (cExN) and inhibitory interneurons (cIN), as disruption of both of these neuronal cell types is frequently implicated in neurodevelopmental disorders. This work defined a suite of affectation-linked neurodevelopmental anomalies in the AP, including deficits in neurite extension, neuronal migration and maturation, and ER stress, and related perturbations of neuronal gene expression, as defined by RNA-seq analysis. While these neurodevelopmental deficits were not present in the UM, the UM and AP models shared multiple functional abnormalities, as defined by electrophysiological analysis, including increased action potential firing and enhanced choline responsiveness.

To our knowledge, this is the first cellular modeling study to characterize cases of *CHRNA7* duplication involving two first degree relatives with differential affectation. A range of molecular, cellular, and functional assays were used to define alterations of neuronal gene expression, neurodevelopment, and altered functional properties linked either to affectation or to *CHRNA7* duplication, both with and without affectation, and some affectation-linked phenotypes were amenable to pharmacological rescue. The distinct neuronal anomalies related to affectation versus *CHRNA7* duplication that were defined here could contribute to the differential phenotypic penetrance seen in other duplication carriers, and provide models that can be used for pre-clinical testing of potential interventions for individuals with this disorder.

## Methods

### Clinical phenotypes

The AP’s pregnancy was planned and there was no history of *in utero* exposure to drugs, alcohol, or tobacco. There were no complications during the pregnancy and the AP was delivered normally at full term, with a birth weight of seven pounds, six ounces. As an infant, the AP was delayed in sleeping through the night, which did not begin to occur until after 1 year of age. He has no history of delay in motor development but has a history of significant delay in language development, producing his first word after the age of two years. He started speech therapy at two years of age, resulting in rapid improvement in his language development. The AP was twelve years old when this study was initiated, with a history of ADHD, depression, and ASD. Prior to age five, he reportedly did not make eye contact and did not exhibit age-appropriate social reciprocity. While in the third grade, the AP had frequent crying episodes and was overwhelmed by homework, which brought him to psychological evaluation. At that time, he manifested both autistic features and pronounced mood lability on exam, which was manifested on several occasions by the child becoming morose and tearful when told even slightly sad stories that would not have elicited such a reaction in a typical child his age. During periods of depressive symptomatology, his mother reported that he had difficulty falling asleep, low appetite, and decreased interest in his favorite activities, such as sports. In middle school, he manifested poor concentration and difficulty with time management, standardized test taking, and organizing tasks and activities. These issues were treated by using cognitive behavioral therapy and play therapy. He was subsequently treated with sertraline, followed by escitalopram. On these selective serotonin reuptake inhibitors, the AP’s anxiety was significantly lessened, but he then experienced residual lack of motivation and his perseverative traits were not improved by treatment. Based on his developmental history, including language delay, impairment in social reciprocity and non-verbal communication, and repetitive thinking, it was clinically determined that a significant contribution to his overall impairment was autistic perseveration and rigidity, for which a trial of risperidone was initiated and resulted in significant clinical improvement over the ensuing years. Ultimately, he was successfully weaned from risperidone and was reasonably well adapted in high school. The AP’s mother reported a history of mild depression, anxiety, and obsessive-compulsive traits, while the AP’s eight-year-old brother had subtle autistic traits that were less pronounced than those of the AP and behavioral features of emotional dysregulation that were more pronounced than those of the AP; he met criteria for disruptive mood dysregulation disorder, ADHD, and mood disorder. Clinical phenotypes of these three subjects with 15q13.3 duplication are summarized in **Table 1**.

### Genotyping

Cytogenetics Microarray (CMA) analysis was performed for research testing by the Washington University Cytogenetics and Molecular (CMP) Pathology Laboratory, using the Affymetrix CytoScan HD array. This array includes 2.6 million copy number markers, 1.9 million non-polymorphic probes, and nearly 750,000 single nucleotide polymorphism (SNP) probes. Average intragenic marker spacing is equivalent to 1 probe per 880 basepairs. Analysis of these data by the CMP Laboratory, after alignment to hg19, defined a 424 kb gain at 15q13.3 in samples from the Affected Proband (AP) and Unaffected Mother (UM) and a 444 kb gain in the same location in the Affected Brother (AB). This CNV was not present in the father.

### iPSC generation

The Washington University Genome Engineering and Induced Pluripotency Center (GEiC) derived multiple clonal iPSC lines from individuals in this family. Briefly, fresh urine samples were procured from the AP and UM and were used to obtain renal epithelial cells, which were reprogrammed using the CytoTune-iPSC 2.0 Sendai virus-based reprogramming kit (Thermo Fischer Scientific), as per the manufacturer’s instructions. iPSC clones were picked and three clonal cell lines were derived from each study subject. Clones number 1 and 3 from each subject were used for experimentation.

### iPSC cultures and differentiation

iPSC lines were grown on Matrigel (Corning) under feeder-free conditions using mTeSR1 (STEMCELL Technologies). For directed differentiation to generate cortical excitatory neural progenitor cells (cExNPCs), iPSCs were dissociated into single cells with Accutase (Life technologies) and 40,000 cells were seeded in V-bottom 96 well non-adherent plates (Corning). Embryoid bodies (EBs) were generated by centrifugation of the plate at 200xg for five minutes, and were then incubated in 5% CO2 at 37°C, in cExNPC differentiation medium with 10μM Y-27632 (Tocris Biosciences). cExNPC differentiation medium includes Neurobasal-A (Life Technologies), 1X B-27 supplement without Vitamin A (Life Technologies), 10μM SB-431542 (Tocris Biosciences), and 100nM LDN-193189 (Tocris Biosciences). On day four, EBs were picked with wide bore P1000 tips and were transferred to Poly-L-Ornithine- (20μg/ml) and laminin- (10μg/ml) coated plates. Every other day media without Y-27632 was replenished and on day 15 Neural Rosette Selection Reagent (STEMCELL Technologies) was used to isolate cExNPCs from rosettes, as per the manufacturer’s instructions. cExNPCs were grown as a monolayer using cExNPC differentiation media up to 15 passages.

Cortical inhibitory neural progenitor cells (cINPCs) were generated by directed differentiation in media which included Neurobasal-A (Life Technologies), 1X B-27 supplement without Vitamin A (Life Technologies), 10μM SB-431542 (Tocris Biosciences), 100nM LDN-193189 (Tocris Biosciences), 1μM Purmorphamine (Calbiochem) and 2μM XAV-939 (Tocris Biosciences). Y-27632 was also included in this media until day eight. For cINPC differentiation, EBs were generated as described above for cExNPC differentiation. At day 4, EBs were transferred to non-adherent plates and were placed on an orbital shaker at 80rpm in an incubator with 5% CO2 and 37°C. cINPC media was replenished every other day and, at day ten, EBs were transferred to Matrigel- and laminin- (5μg/ml) coated plates. On day 15, cINPCs were dissociated with Accutase and were either cryopreserved and/or grown in monolayer culture on Matrigel- and laminin-coated for up to 15 passages. For analysis of both cExNPC and cINPC growth properties, equal numbers of cells for each line were seeded on Matrigel- and laminin- (5ug/ml) coated plates and the total number of cells was counted after four days.

For differentiation and maturation of neurons, cortical neuroids were generated by seeding both 2X10^4^ cExNPCs and 2X10^4^ cINPCs into each well of a V-bottom 96 well non-adherent plate in maturation media. Plates were spun at 200xg for five minutes and were incubated in 5% CO2 at 37°C in maturation media, with addition of Y-27632 for the first four days of culture. The composition of maturation media includes Neurobasal-A and 1X B-27 supplement without Vitamin A, while DAPT (10μM; Tocris) were included in the media from day 7 to day 11, and 200μM Ascorbic acid (Sigma Aldrich), 20μg/ml BDNF (Peprotech), and 200μM cAMP were included in the media from day 11 to day 15. At day 4, neuroids were transferred to non-adherent plates and were placed on an orbital shaker at 80rpm in an incubator with 5% CO2 and 37°C. At day 5, neuroids were moved to Matrigel- and laminin- (5μg/ml) coated plates and further incubated in 5% CO2 at 37°C. Media were replenished every other day until day 15.

### Immunocytochemistry (ICC) and Immunoblotting

ICC and immunoblotting experiments was performed as previously described (22). In brief, for ICC, cortical neuroids were dissociated after 15 days of maturation and cells were plated in eight-well chamber slides coated with Matrigel and laminin (5μg/ml). After 24 hours, cells were washed with PBS without calcium and magnesium and were fixed in 4% paraformaldehyde for 15-20 minutes. See (22) for detailed protocol. Primary and secondary antibodies used for these experiments are provided in **Supplemental Table S1A**. Images were taken using a spinning-disk confocal microscope (Quorum) and an Olympus inverted microscope using MetaMorph software. ImageJ was used to process images and for quantification: 15-20 random fields were imaged from three to five biological replicate experiments which included work with two different clones per subject; total numbers of both immune-positive and all DAPI stained cell nuclei quantified are shown in **Supplemental Table S1B**.

### FACS analysis

Approximately one million cExNPCs or cINPCs between passages 4-9 were used for FACS analysis and experiments were performed as described previously (22). P-values: * P<0.05, **P<0.01, ***P<0.001 were determined by unpaired t-test.

### RNA-Sequencing and RT-qPCR

After 15 days of cortical neuroid differentiation as described above, total RNA was collected from the AP, UM, and Unrelated Control Male (UC-M) and Unrelated Control Female (UC-F) lines, using the NucleoSpin RNA II kit (Takara) per the manufacturer’s instructions. RNA was quantified using a NanoDrop ND-1000 spectrophotometer (Thermo Scientific) and the Agilent Bioanalyzer 2100 was used to assess RNA integrity, with only samples with an RNA Integrity Number of >8 used for sequencing and analysis. RNA-Sequencing (RNA-Seq) library preparation and Illumina Sequencing were performed by the Genome Technology Access Center at Washington University. The Illumina Hi-Seq3000 was used to obtain single end 50 base pair reads, with approximately 30 million unique reads per sample obtained after alignment. Four independent biological replicates per cell line were analyzed by RNA-Seq. For RT-qPCR, 1μg total RNA was reverse transcribed using iScript Reverse Transcription Supermix (Bio-Rad) and equal quantities of cDNA were used as a template for RT-qPCR using the Applied Biosystems Fast Real Time quantitative PCR platform. GAPDH or RPL30 were used as endogenous controls for normalization. Four biological replicate experiments using one clonal line for each sample type were used for RNA-seq analysis (n=4), while a second clonal line for the UM and AP models was used to generate RNA for RT-qPCR validation of a subset of the RNA-Seq findings. P-values for RT-qPCR validation: * P<0.05, **P<0.01, ***P<0.001 were determined by unpaired t-testing.

### Bioinformatics and IPA analysis

RNA-Seq data analysis was performed as described in (22, 32) to obtain differentially regulated genes (DEG). Briefly, STAR version 2.5.4b was used to align the RNA-Seq reads to the human genome assembly hg38 (22). To derive uniquely aligned unambiguous reads, we used Subread:featureCount, version 1.6.3 with GENCODE gene annotation (22) and gene-level transcripts were imported into the R-bioconductor package (22). After excluding genes expressed at <1.0 counts per million (CPM), Differentially Expressed Genes (DEG) were curated based upon a Log2 fold change >1 and a Benjamini and Hochberg FDR of <0.05. DEGs were used to perform hierarchical clustering analysis using ClustVis (33) and to perform Ingenuity Pathway Analysis (IPA) (Qiagen), as described previously (22). To determine the contribution of different covariates to gene expression, we also performed variancePartition analysis and including the individual sample types (UC-M, UC-F, UM and AP), age (young and old), and sex (male and female) as variables (34).

### Morphometric analysis

To measure neurite extension, cortical neuroids were generated and cultured to promote differentiation and maturation as described above. On day 6, images were acquired using an inverted light microscope for the UC-M, UM, and AP samples, with data collected for three or more independent biological replicate experiments encompassing work with two clonal lines per subject for the AP and UM, and in one clonal line for the UC-M. For each experimental finding in this manuscript, the number of biological replicate experiments and the clones used for each replicate of each type of experiment are summarized in Supplemental Table S1C. Neurite extension length from adherently plated neuroids was measured using ImageJ, as the distance between two circles drawn at the border of the plated neuroid and at the tips of neurites extending from that neuroid, as shown in **Fig. 3A**. ICC for MAP2 in adherent neuroids was conducted as described above, with image acquisition using a spinningdisk (Quorum) confocal microscope and an Axiovision inverted microscope. Day 15 neuroids were also dissociated and the neurons plated on Matrigel- and laminin-coated plated and stained with MAP2. To assess neurite length, images were acquired using a spinning-disk (Quorum) confocal microscope and an Axiovision inverted microscope. Images were processed with Imaris software (Bitplane) and neurite length was measured using the filament tracer application and normalized to the number of nuclei stained with DAPI in the neuroid, which was measured with particle application in Imaris. To quantify VGAT- and VGLUT-expressing punctae, dissociated and plated neuroids generated as described above were immunostained for the respective antibodies, and images taken using a spinning-disk (Quorum) confocal microscope and Axiovision inverted microscope. Punctae were measured using the synaptic counter plugin in ImageJ. Each finding was obtained in three or more independent biological replicate experiments encompassing work with two clonal lines per subject for the AP and UM, and one clonal line for the UC-M.

### Migration assay

To study migration of cExN and cIN neurons, we developed an approach that utilized fused co-culture of two 3D spheres consisting of cExNs and cINs. These 3D spheres were generated by transducing cExNPCs and cINPCs separately with either a lentiviral synapsin-eGFP or a synapsin-RFP expression construct, respectively. 30,000 of these transduced cEXNPCs or cINPCs per well were then seeded into separate wells of a V bottom 96 well plate in 100μl of maturation media containing 10μM Y-27632. The V bottom plate was centrifuged at 200xg for 5 minutes at room temperature and then incubated in 5% CO2 at 37°C. On day 2, 50μl of media was replaced with fresh media without disturbing the spheres. On day 4, cEXN and cIN EBs were selected with wide bore P1000 tips and moved to a U bottom plate. One cExN and one cIN sphere were placed side by side in each U bottom well. Placement of spheres in close apposition caused them to undergo fusion without further manipulation, enabling assessment of neuronal migration. On day 6, these fused spheres were moved with wide bore P1000 tips to a coverslip placed in a 3cm plate and coated with matrigel and laminin (5μg/ml). On day 10, images were acquired using a spinning-disk (Quorum) confocal microscope and Axiovision inverted microscope and image processing was performed using ImageJ. Migration was assessed in such fused co-cultures derived from the UC-M, UM, and AP lines. The ability of WNT signaling to rescue migration deficits observed in the AP line was tested by addition of 10μM CHIR-99021 in DMSO, with parallel treatment of control spheres with equal quantities of DMSO. Three or more independent biological replicate experiments were performed in two clonal lines for the AP and UM, and in one clonal line for the UC-M. P-values: * P<0.05, **P<0.01, ***P<0.001 were determined by unpaired t-test.

### ER stress luciferase assay

To test the effects of endoplasmic reticulum (ER) stress on cINPCs, we used an expression construct encoding a stress sensor (35). This construct encodes a Guassia luciferase protein fusion, with replacement of the first 18 amino acids with the signal peptide from the mesencephalic astrocyte-derived neurotrophic factor (MANF) protein and carboxy-terminal fusion to MANF’s final 5 amino acids, which encode a stress sensor. To detect ER stress, 35,000 cINPCs were seeded on Matrigel- and laminin- (5μg/ml) coated 96 well plates in cIN differentiation media containing Y-27632. After 24 hours, the cells were transfected with the stress sensor expression construct using FuGENE6 (Promega) transfection reagent. 48 hours after transfection, 50μl of supernatant was removed and assayed for luciferase activity using the BioLux Gaussia Luciferase Assay Detection System (New England Biolabs). For rescue experiments, small molecules were obtained from Sigma Alrdich or Tocris Biosciences, reconstituted in DMSO or PBS-Ca^2+^/Mg^2+^, and added in the medium after 24 hours of transfection, at the final concentrations indicated (Tudca-50μM, PBA-500μM, Dantrolene sodium-1μM, and JTV-519-10μM). Luciferase levels were measured after 48 hours of small molecule treatment as above. Luciferase data for rescue experiments performed in presence of small molecules were normalized to the DMSO control, and three or more independent biological replicate experiments were performed, using two clonal lines for the AP and UM and one clonal line for the UC.

### Electrophysiology

cExNPCs and cINPCs were transduced with Synapsin-GFP and Synapsin-RFP expression constructs, respectively, before performing cortical neuroid maturation. At day 0 of cortical neuroid differentiation, 2X10^4^ cExNPCs and 2X10^4^ cINPCs were mixed in each well of a V-bottom 96-well non-adherent plate in maturation media. Maturation followed the approach described above. At day 15, neuroids were dissociated using Accutase (Life Technologies), were seeded onto a layer of rat cortical astrocytes, prepared as described previously (32), and were grown for another three weeks using Neurobasal-A, 1XB27 with vitamin A (Life Technologies), and supplementation with BDNF (20ng), cAMP (200μM) and ascorbic acid (200μM). iPSC-derived neuron cultures were perfused at 1 ml/min with Tyrode’s solution (in mM): 150 NaCl, 4 KCl, 2 MgCl2, 2 CaCl2, 10 Glucose, 10 HEPES, with pH adjusted to pH 7.4 with NaOH. Recording electrodes had an open-tip resistance of 2-6 MOhm when filled with (in mM): 140 K-glucuronate, 10 NaCl, 5 MgCl2, 0.2 EGTA, 5 Na-ATP, 1 Na-GTP, and 10 HEPES, pH adjusted to 7.4 with KOH. Whole-cell currents and membrane potentials were recorded with an Axopatch 200A amplifier (Molecular Devices). Voltage clamp recordings were used to determine cell capacitance and input resistance as well as peak inward sodium current and steady-state outward potassium currents during depolarizing voltage steps from a holding potential of −80 mV (32). Current clamp recordings and currents evoked by choline and acetylcholine (ACh) were obtained in a modified extracellular solution (in mM): 120 NaCl, 3 KCl, 10 glucose, 1 NaH2PO4, 4 NaHCO3, 5 HEPES, pH adjusted to 7.4 with NaOH, delivered from an 8-barrelled local perfusion pipette positioned near the recorded cell.

## Results

### iPSC-derived neural progenitor cells from the affected proband exhibit increased proliferation

The pedigree selected for study includes three individuals carrying 15q13.3 duplication: the mother, who is not clinically affected (UM), her older son, who exhibits distinct features of autism and emotional dysregulation (the affected proband, AP), and her younger son, who exhibits mild ASD, ADHD, and mood disorder traits (the affected brother, AB). Chromosomal Microarray Analysis (CMA) conducted on the AP, UM and AB, defined one CNV in these individuals, an approximately 400 kilobase gain at chromosome 15, band q13.3 (see Methods, **Table 1**, **Supplemental Figure S1**). Only one gene, *CHRNA7*, is located within the duplicated region. The father in this pedigree does not carry this duplication (**Supplemental Figure S1**). Clinical phenotypes of the three mutation carriers are summarized in **Table 1**.

Renal epithelial cells isolated from the UM and AP (**Sup. Figure S1**) were used for reprogramming to generate three clonal iPSC lines from each of these individuals, and these lines were compared with single clonal iPSC lines derived from unrelated, unaffected male and female control subjects (UC-M/UC-F). All lines exhibited a similar morphology, expressed the pluripotency marker OCT4 (POU5F1), and were karyotypically normal (**Sup. Figure S2**). The development and/or function of cortical excitatory neurons (cExNs) and inhibitory cortical interneurons (cINs) is frequently disrupted to contribute to neurodevelopmental disorders (36, 37). Therefore, we used two clonal lines each from the UM and AP, and one clonal line each from the UC-M and UC-F to generate cExN and cIN neural progenitor cells (cExNPCs and cINPCs) (**Figure 1A**). After 15 days of NPC specification, cEXNPCs and cINPCs were maintained as a monolayer as described in the methods. CHRNA7 expression levels were assessed by RT-qPCR and were significantly increased in both the AP- and UM-derived NPCs, by comparison with the UC-M-derived NPCs (**Figure 1B**). During cExNPC and cINPC maintenance, the AP-derived NPCs appeared to exhibit more rapid proliferation than the UM- or UC-M-derived NPCs, as measured by seeding equal numbers of cells and quantitation after four days (**Figure 1C, 1F**). We further assessed this by performing FACS analysis of propidium iodide (PI)-stained cExNPCs and cINPCs, finding that the AP NPCs had significantly higher percentages of both S- and M-phase cells than either the UM or UC-M NPCs. Representative FACS plots and summary data, indicating percentages of cells in the S and 4N phases of the cell cycle are shown for cExNPCs and cINPCs, respectively (**Figure 1D-E, 1G-H**), with summary data for all cell cycle phases shown in **Supplemental Fig. S3**, and numbers of biological replicate experiments and clones used for each experiment in this manuscript summarized in **Supplemental Table S1C**.

**Figure 1.**
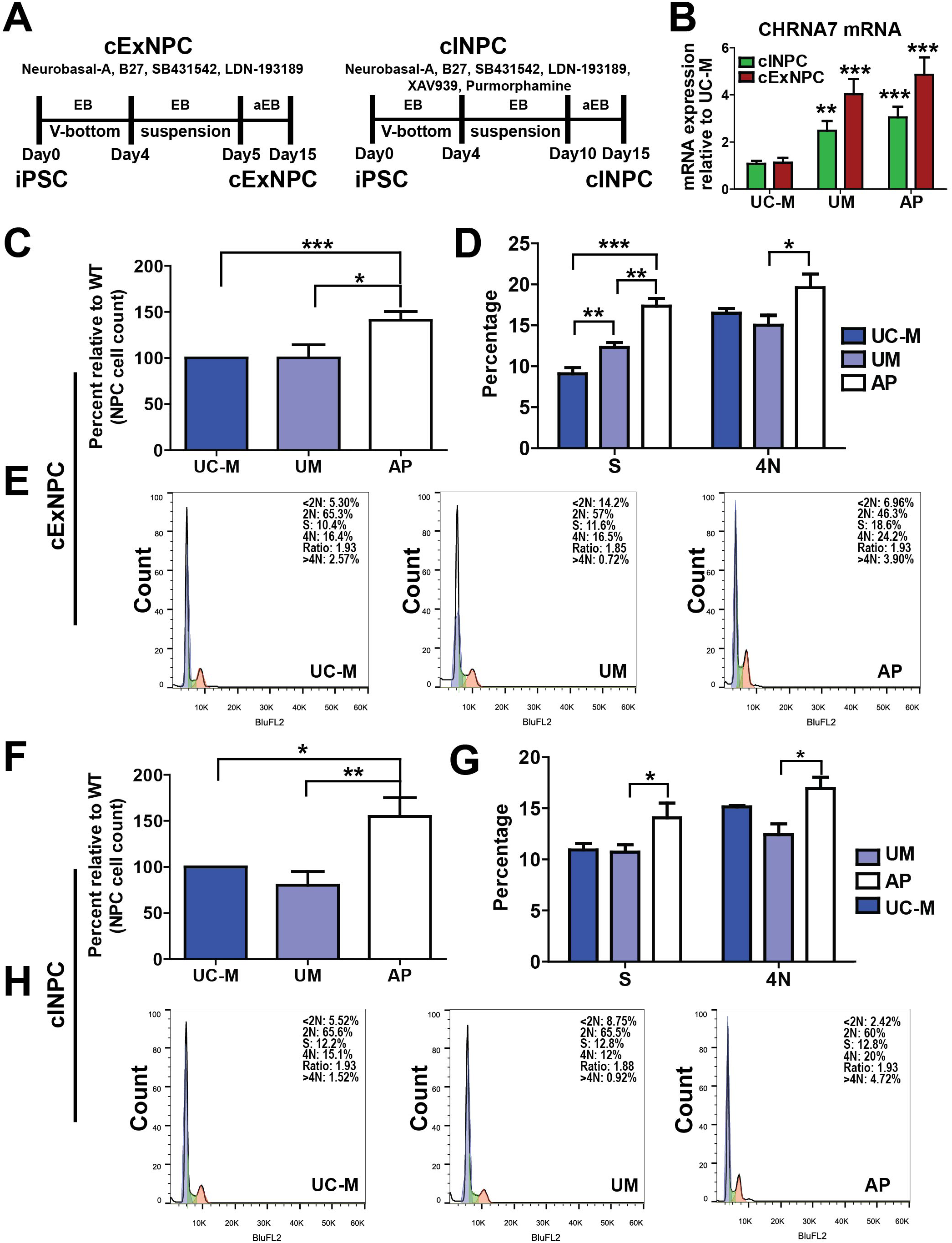
Generation and characterization of cExNPCs and cINPCs from patient-derived iPSCs. (A) Differentiation schemes used for generation of cExNPCs and cINPCs, including timeline and small molecules used and maintenance as a suspended embryoid body (EB) or adherent EB (aEB). (B) CHRNA7 expression in cExNPCs and cINPCs was analysed by RT-qPCR. Data shown is the average +/-SEM of results from four independent biological replicate experiments (n=4). (C) cExNPCs were seeded in equal numbers for each line tested and cell numbers were counted after four days of maintenance (n=7 biological replicates). (D-E) cExNPCs were stained with propium iodide for DNA content and analyzed by FACS. S and 4N (M phase) cells were quantified (D) for each study subject. In D, average values are shown from seven independent biological replicates (n=7); representative FACS plots are shown in (E). (F) cINPCs were seeded in equal numbers for each line tested and cell numbers counted after four days of maintenance (n=7 biological replicates). (G-H) cINPCs were stained with propidium iodide for DNA content and analyzed by FACS. S and 4N (M phase) cells were quantified for each study subject. In G, values are shown from seven independent biological replicate experiments (n=7); representative FACS plots are shown in (H). All significant findings in this manuscript were confirmed in three or more independent biological replicate experiments performed using two clonal lines for the UM and AP models, and one clonal line for the UC-M model, as summarized in **Supplemental Table S1C**. p-values *\*P* < 0.05, *\*\*P* < 0.01, *\*\*\*P* < 0.001 were determined by unpaired t-test.

### Defects in neurite extension and production of VGAT-expressing punctae in AP-derived neurons

To assess the differentiation potential of NPCs derived from the UC-M, UM, and AP lines, cExNPCs and cINPCs were combined at a 1:1 ratio to generate cortical neuroids (**Figure 2A**). We used this co-culture approach to accelerate neuronal differentiation and maturation, and to model interactions between cortical excitatory and inhibitory neurons that can occur during cortical development *in vivo*. Upon dissociation and plating of these day 15 neuroids on 8 well chamber slides, AP-derived neuroids exhibited increased expression of NPC markers including DLX2, TBR1, and PAX6, relative to the UM and UC neuroids, while neuroids also contained higher frequencies of cells expressing the proliferative marker Ki67 (**Figure 2B-F**). This could be related to the increased proliferation of the AP-derived NPCs, relative to the UM- and UC-derived NPCs (**Figure 1**). We also plated these neuroids for differentiation without dissociating them, allowing them to extend neurites. By 5 days of differentiation, the AP-derived neuroids exhibited a deficit in neurite extension not seen in either the UC- or UM-derived neuroids. This reduction in neurite extension was quantified using light microscopy and further visualized by MAP2 staining of the day 5 plated neuroids (**Figure 3A-C**). On day 15, neuroids were dissociated and the plated neurons were used for ICC analysis. MAP2 staining of these neurons confirmed a neurite length deficit in the AP-derived neurons (**Fig. 3D-3D’**). ICC for the GABA and glutamate transporters VGAT and VGLUT was further used to assess formation and transport of synaptic vesicles in differentiatied cINs and cExNs, respectively. While VGLUT-expressing punctae were present in cExN neurites derived from all three lines, AP-derived cINs exhibited significantly reduced formation of VGAT-expressing punctae along neurites, compared to those derived from the UC-M and UM (**Figure 3E-E”**). Both the increased expression of NPC and proliferative markers and the impaired acquisition of characteristics of differentiated and mature neurons suggested that the AP-derived neurons were relatively immature, by comparison with those derived from the UM and UC-M.

**Figure 2.**
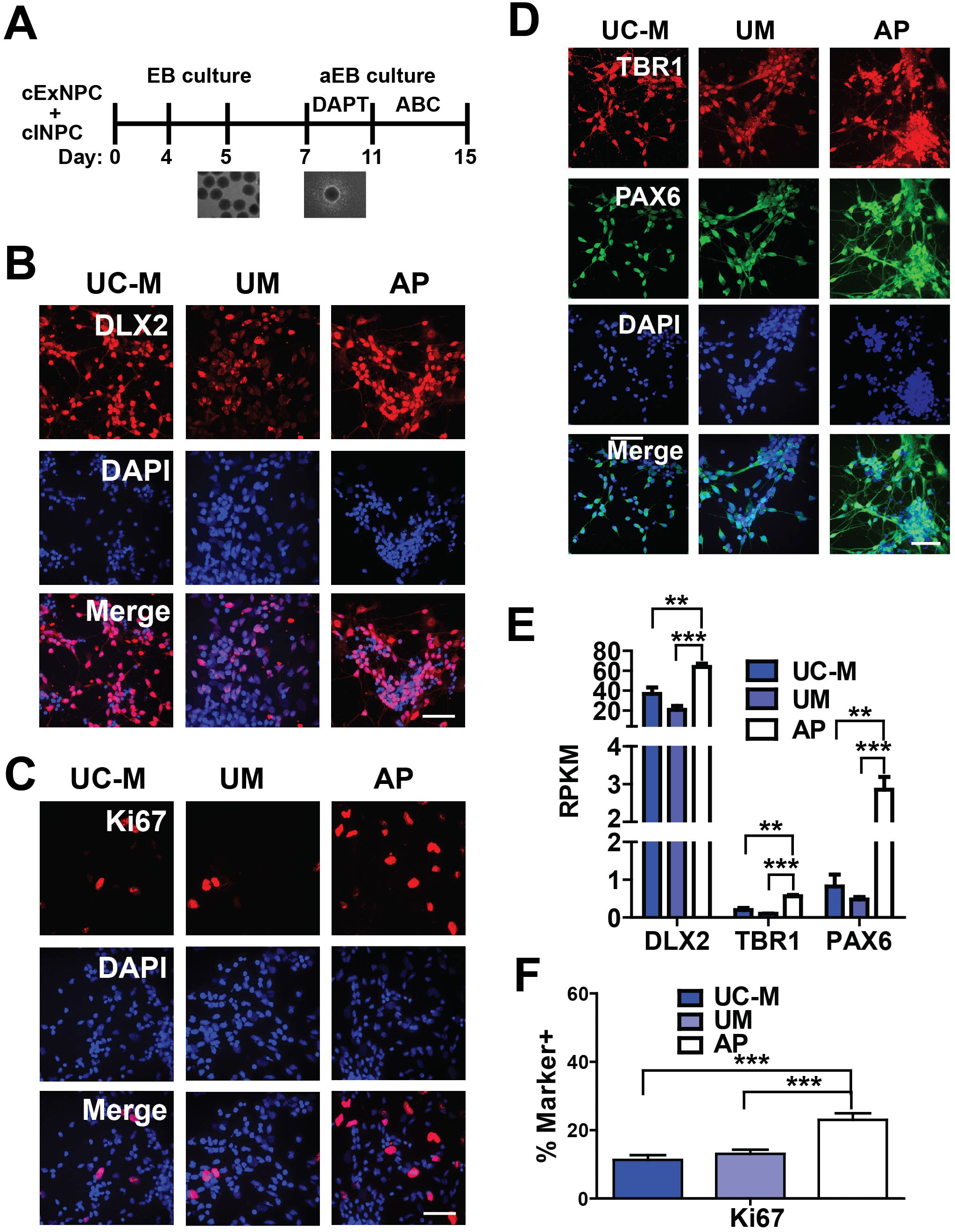
Generation and characterization of cortical neuroids. (A) Schematic of the method used to generate cortical neuroids, by combining cExNPCs and cINPCs in a 1:1 ratio and differentiating and maturing them for fifteen days. ABC=Ascorbic acid, BDNF, and cAMP. See Methods for further details. (B-D) Immunocytochemistry with the antibodies indicated detects cINPCs (DLX2), proliferating NPCs (Ki67), and cExNPCs (TBR1 and PAX6), with representative images from one clonal line per subject shown. (E) RNA-seq analysis defined differences in gene expression between the AP, UM, and UC-M neuroids for the markers shown (n=4 independent biological replicates from one clonal line per subject). (F) Immunocytochemical quantification of the percentage of cells expressing the proliferative marker Ki67, n=4 biological replicate experiments utilizing two clonal lines for the AP and UM and one clonal line for the UC-M. Scale bars=50μm.

**Figure 3.**
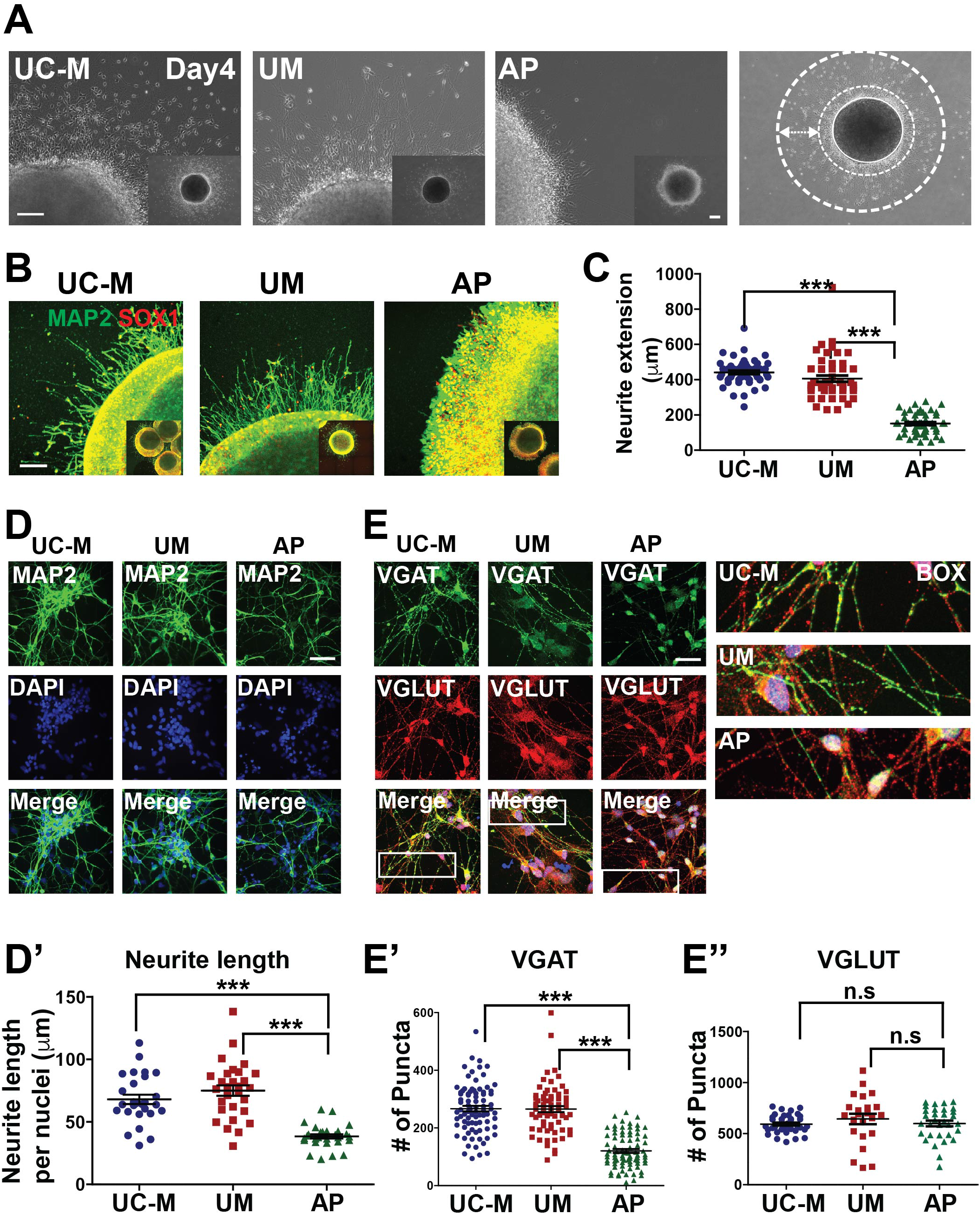
Morphometric analysis of differentiated cortical neuroids. (A-C) Five days after plating cortical neuroids in differentiation media, protruding neurites were analyzed for each sample type. (A) Representative light microscopy images are shown for each sample type. Quantification was performed as shown on right, by defining the distance between two circles drawn at the border of the plated neuroid and at the tips of neurites extending from that neuroid. Scale bar=250 μm. (B) Immunocytochemical analysis of neurite extension using MAP2 staining, with representative images shown. Scale bar=150 μm. (C) Quantification of neurite extension is shown for seven biological replicate experiments (n=7). (D-D’) Neurite length was analyzed in plated MAP2 immunostained neurons derived from cortical neuroids. Representative images are shown in (D) and neurite length is quantified in D’, using data from three independent biological replicate experiments (n=3). (E-E’’) Expression of the GABA and glutamate transporters, VGAT and VGLUT, was assessed by immunocytochemical analysis of neuroids. Representative images are shown in (E) and synaptic puntae were quantified in (E’-E’’) for VGAT (E’) and VGLUT (E’’). Data were derived by quantification of 15 stained neuroids derived from four independent biological replicate experiments (n=4). Scale bar=75 μm. p-values *\*P* < 0.05, \*\**P* < 0.01, \*\*\**P* < 0.001 were determined by unpaired t-test.

### Transcriptome comparisons in cortical neuroids

To identify differences in gene expression that distinguished neurons derived from the AP, UM, UC-M, and UC-F lines, we performed RNA-seq analysis on cortical neuroids after 15 days of neuronal differentiation. Four independent biological replicate data sets were generated and these were clustered by Principal Component Analysis (PCA) of processed reads (**Figure 4A**). We initially assessed differentially expressed genes (DEGs) specific to the AP, which could be related to the altered differentiation observed in this model, by performing pairwise comparisons with the other sample types (UM, UC-M, UC-F), using cut-off values of FDR <0.05 and log2 fold difference >1 (see Methods, **Supplemental Table S2**). We focused first on genes that were differentially expressed in the AP versus (vs) the UM, as these subjects have a shared genetic background including *CHRNA7* duplication, but exhibit differential affectation (**Fig. 4B**, blue). Hierarchical clustering of these DEGs, with comparisons to the UC samples included for a full sample comparison, demonstrated that the majority exhibited increased expression in the AP, relative to the UM (**Fig. 4C**). We used Ingenuity Pathway Analysis (IPA) to define pathways and disease-related networks enriched among these AP-specific DEG’s (**Supplemental Table S3**). These included molecular pathways involved in axon guidance signaling, the cell cycle, WNT signaling, GABA receptor activity, neuroinflammation, and gap junction signaling (**Figure 4D; Sup. Table S3**), with most genes related to WNT signaling exhibiting diminished expression in the AP, compared with the UM (**Figure 4E**). IPA disease network analysis also defined clusters of genes with differential expression in the AP related to nervous system development, developmental disorders, behavior, and psychological disorders (**Figure 4F**). For example, clusters of genes related to the disease terms cognition and neuritogenesis exhibited predominantly reduced expression in the AP-derived neurons (**Figure 4G-H**). The finding that neuritogenesis-related genes were diminished in the AP was congruent with the AP-specific neurite extension defect observed in **Fig. 3A**.

**Figure 4.**
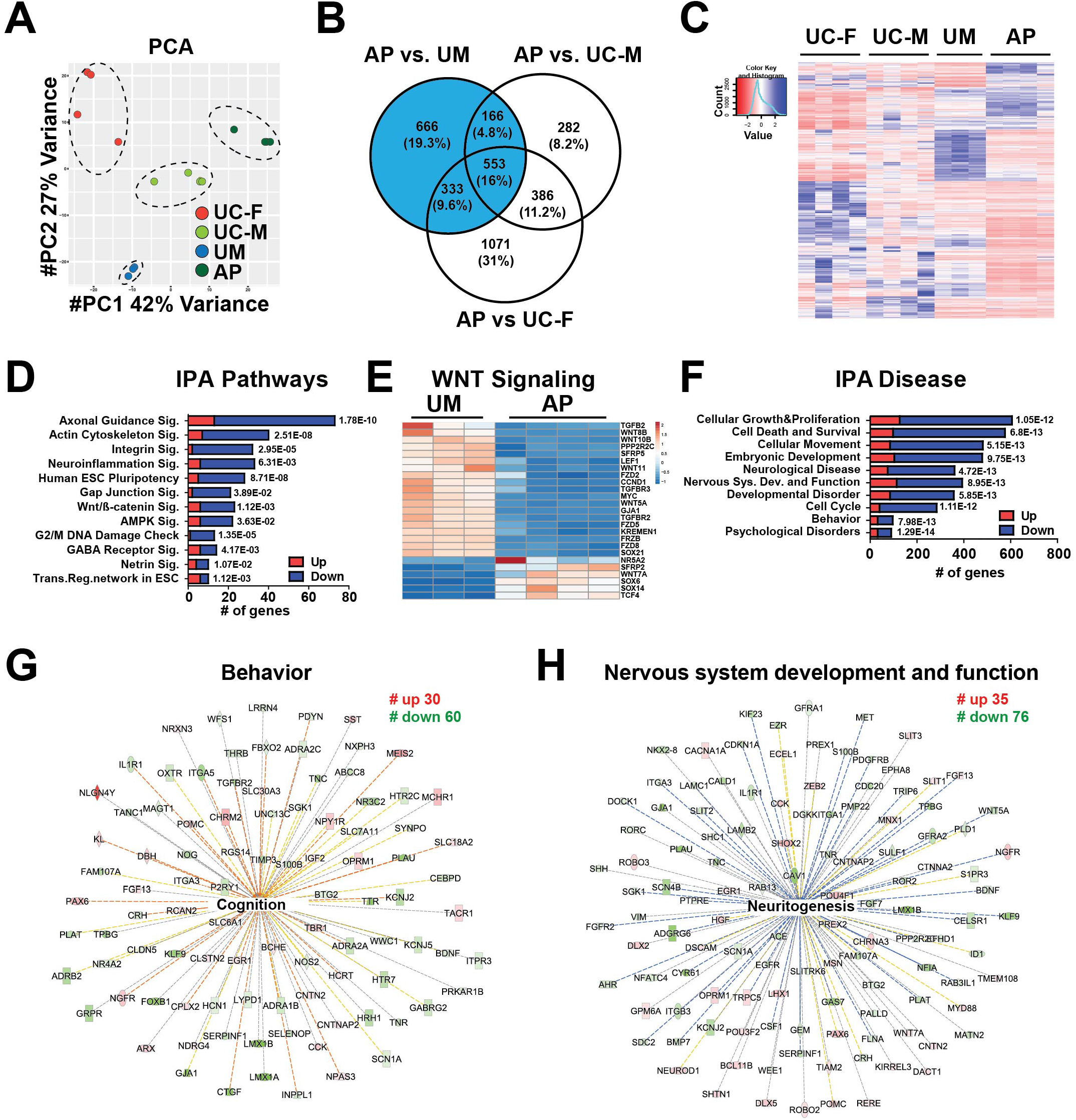
Transcriptomic analysis of differentiated cortical neuroids, defining differential gene expression between the AP and UM. RNA-seq was conducted on cortical neuroids that had been differentiated for 15 days. (A) Principal Component Analysis (PCA) of gene expression in differentiated neuroids derived from the UC-M, UC-F, UM, and AP lines is displayed as a multidimensional scaling plot derived from four independent biological replicate experiments (n=4) performed using one clonal line for each subject. (B) Venn diagram showing differentially expressed genes (DEGs) defined by pairwise comparisons of the AP versus (vs.) UM, AP vs. UC-M, and AP vs. UC-F datasets. AP vs. UM-specific DEGs are shown in blue. (C) These AP-specific DEGs were further analyzed by hierarchical clustering analysis, with other samples included for comparison. (D-H) DEGs were assessed by Ingenuity Pathway Analysis (IPA), identifying (D-E) AP-enriched pathways and (F) disease-related GO terms. (D) IPA-pathway analysis identified differential enrichment for WNT signaling-related gene expression, with expression of these genes visualized as a heat map in (E). In D and F, the number of genes related to each term is represented on the x-axis, while red and blue color indicates up and down-regulated genes, respectively. p-values for each term are indicated to the right of each bar. (F) IPA disease GO terms identified gene networks associated with (G) Behavior and (H) Nervous system development and function. Numbers (#) of up- and down-regulated genes present each network are indicated. Within each network, red and green symbols indicate up- and down-regulated genes, respectively, while color intensity indicates the relative degree of differential expression.

We also curated the DEGs that were AP-specific by comparison with both the UM and unrelated male and female controls lacking the 15q13.3 duplication (UC-M/UC-F)(1052 genes; **Supplemental Fig. S4A**). These included many genes with diminished expression in the AP relative to the other sample types (**Supplemental Fig. S4B**) and encompassed many molecular pathways and disease-related GO terms similar to those identified in the AP versus UM comparison, including WNT signaling, axon guidance signaling, GABA receptor signaling, nervous system development, and psychological disorders (**Supplemental Fig. S4C-F; Sup. Table S4**). For example, clusters of genes related to the growth of neurites and migration of neurons were predicted to have predominantly reduced expression in the AP relative to the other sample types (**Sup. Fig. S4C-F; Sup. Table S4**). This suggested that similar pathways exhibited impaired expression in the AP, by comparison with both the UM and the models derived from unrelated individuals.

### UM-specific differential gene expression in cortical neuroids compared to unrelated controls

Since both related models derived from the AP and UM carry the same *CHRNA7* duplication, we also defined DEGs specific to the UM, by comparison with the male and female unrelated controls (**Supplemental Figure S5A**). A unique cluster of DEGs exhibited increased expression, relative to all other samples, while a cluster with decreased expression relative to the UC-M and UC-F samples was shared by both the UM and AP models (**Supplemental Figure S5B**). IPA analysis of these UM-specific DEGs (**Supplemental Table S5**) indicated enrichment for some gene ontology (GO) terms that were similar to those identified for the AP-specific DEGs (e.g. GABA receptor signaling, gap junction signaling, and behavior/psychological disorders; **Sup. Figure S5C-D**). However, many DEGs and enriched GO terms and pathways obtained from comparisons to the UC models differed between the UM and AP. For example, genes related to WNT signaling were not differentially expressed in the UM vs UC comparisons. Furthermore, while the UM-specific DEGs also included a neuritogenesis-related network, genes in this network were more highly expressed in the UM than in the unrelated controls (**Sup. Figure S5E-F**), while neuritogenesis-related genes had diminished expression in the AP, by comparison with both related and unrelated controls (UM and UC) (**Fig. 4H, Sup. Fig. S4F**). We used cortical neuroids generated using a second clonal line for each of the AP and UM models to confirm a subset of these findings by RT-qPCR, testing integrin molecules, channel related genes, and transcription factors. These analyses demonstated differences in gene expression between the AP and UM models that were similar to those obtained by RNA-seq analysis (**Sup. Figure S6A-B**). To further assess major contributors to differential expression in the AP vs UM or the AP vs UM and UC models, we performed variancePartition analysis. This analysis indicated that the individuals from whom the samples were derived were the major contributor to the differential expression profiles obtained, while the age and sex of the study subjects were minor contributors (**Supplemental Figure S6C**).

### Interneuron migration is diminished in AP-derived neurons and this defect is partially rescued by a WNT agonist

Analysis of the AP-specific DEGs identified a group of genes that regulate migration of neurons with diminished expression in the AP, by comparison with the UM (**Fig. 5A**). Therefore, we hypothesized that the AP model may have altered neuronal migration capacity. *In vivo*, cortical interneurons undergo tangential migration from the ventral telencephalon to the cortex. Therefore, we adapted an organoid-based model to assess potential alterations of neuronal migration, as described in the Methods. A neuroid consisting of synapsin-GFP expressing cExNs was apposed with a neuroid consisting of synapsin-RFP expressing cINs, and migration of neurons from one neuroid to the other was evaluated (**Figure 5B**). In this assay, cINs from AP neuroids exhibited reduced migration by comparison with both the UM and UC-M (**Figure 5C-5D**). By contrast, the AP cExN neuroids exhibited increased migration by comparison with the UM but not the UC-M. The UM-derived cExNs also exhibited diminished migration by comparison with the UC-M control, suggesting that their migration was impaired (**Figure 5C, 5E**). As WNT signaling has been linked to neuronal migration (38), we hypothesized that the diminished expression of genes involved in WNT signaling in the AP (**Figure 4E**) could contribute to the reduced cIN migration observed. Indeed, treatment of AP neuroids with the WNT agonist CHIR-99021 enhanced cIN migration in the AP model, partially rescuing this impaired migration (**Figure 5C-D**).

**Figure 5.**
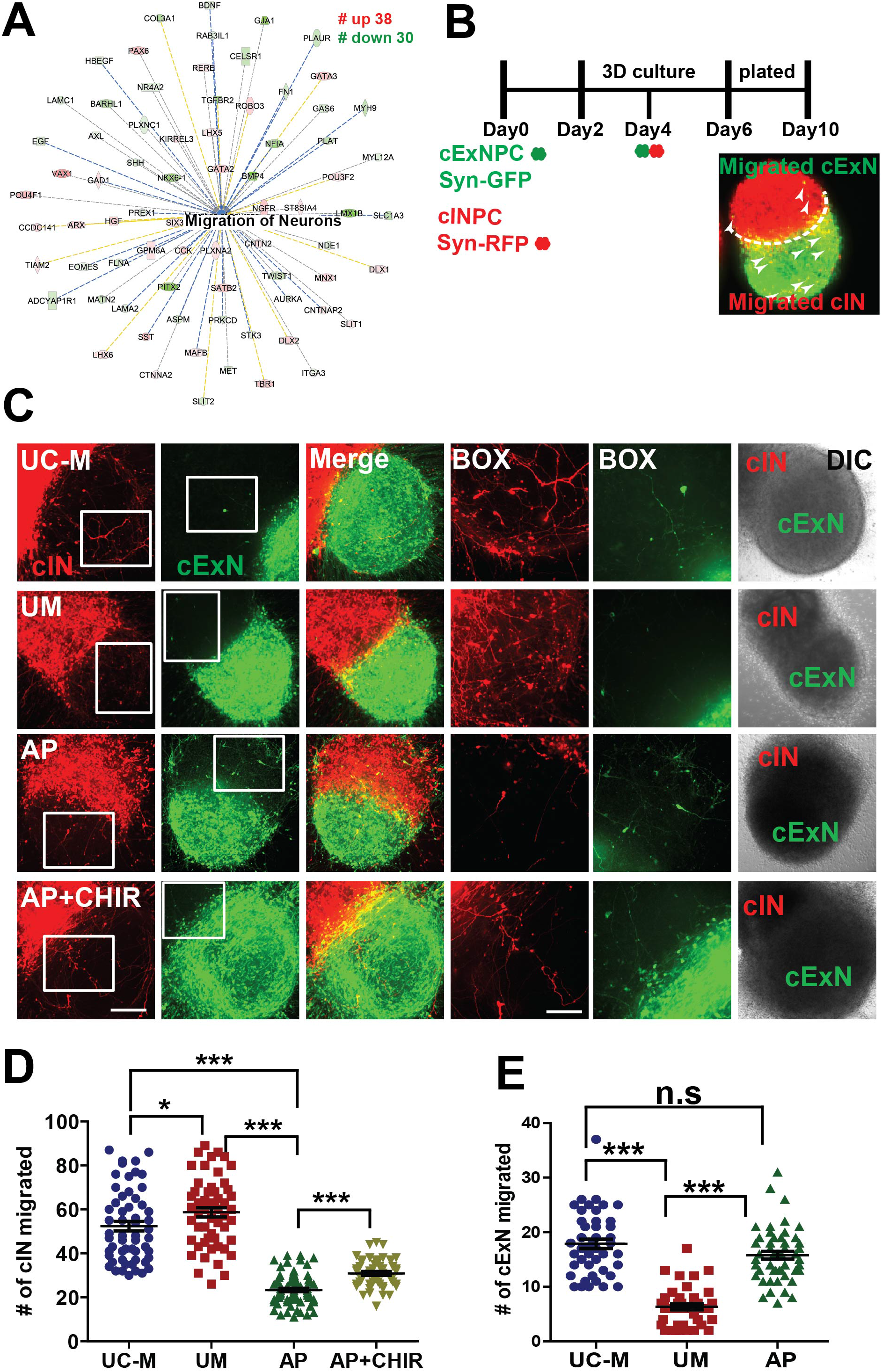
Neuronal migration is compromised in AP-derived cINs and this phenotype is partially rescued by the WNT agonist CHIR-99021. (A) IPA analysis of AP-enriched DEGs (versus the UM sample, Fig. 4 above) identified a cluster of genes which control neuronal migration. (B) Schematic depicting the migration assay, which involves generation of neuroids containing Synapsin promoter (Syn)-GFP-expressing cExNPCs (green) and Syn-RFP-expressing (red) cINPCs, apposition of these neuroids, and differentiation and migration of neurons in these co-cultures, with analysis at Day 10. Neurons that migrated into the opposite neuroid are indicated by white arrowheads. (C) Migration of red cINs into the green cExN neuroid, and vice versa, is shown in representative confocal images from assays performed with UC-M, UM, and AP model-derived neuroid cocultures. (D) The number of cINs (red) that migrated into the cExN neuroid were quantified from six independent biological replicate experiments (n=6), that used two clonal lines for the UM and AP models and one clonal line for the UC-M model. The reduced cIN migration in the AP model was partially rescued by addition of CHIR-99021 (CHIR). (E) The number of cExNs (green) that migrated into the cIN neuroid were quantified, using data from six independent biological replicate experiments (n=6) that used two clonal lines for the UM and AP models and one clonal line for the UC-M model. Scale bars=150 μm and higher magnification (BOX)=100 μm. p-values \**P* < 0.05, \*\**P* < 0.01, \*\*\**P* < 0.001 were determined by unpaired t-test.

### Increased ER stress selective to the AP-derived NPCs was rescued by the ryanodine receptor antagonist JTV-519

In a prior study, a microduplication at chromosome 15q13.3 increased mRNA expression levels of *CHRNA7* and also resulted in moderately increased expression of several endoplasmic reticulum (ER) chaperone and unfolded protein response (UPR)-related ER stress markers (31). Therefore, we assessed whether these models exhibited altered expression of ER chaperones in neuroids, but did not observe increased expression of these or other ER chaperone or UPR-related genes in AP samples, by comparison with UM and UC-M model-derived samples (**Figure 6A**; not obtained as DEGs in **Supplemental Table S2**). However, we found that the brain expressed ryanodine receptor, RYR3, exhibited increased expression in the AP by comparison with the UM and UC samples (**Figure 6B**). As ryanodine receptors modulate calcium homeostasis following increases in ER stress, we hypothesized that the AP could have elevated ER stress due to altered calcium homeostasis, rather than elevated ER chaperone or UPR pathway activities. To assess whether any of these models exhibited altered ER stress, we introduced an expression construct encoding a Secreted ER Calcium Monitoring Protein (SERCaMP)-luciferase stress sensor (35) into cINPCs, using this sensor to monitor calcium release from the ER in response to stress, as detailed in the Methods. Interestingly, cINPCs from the AP selectively exhibited increased ER stress in these assays, while levels seen in the other model with 15q13.3 duplication (the UM) did not differ from that of unrelated controls (**Figure 6C**). As ER stress both activates the UPR and triggers calcium release through ryanodine receptor release channels, we tested whether chemical antagonists of either of these processes could rescue the AP model’s elevated ER stress response. While neither antagonist of the UPR (PBA/TUDCA) exhibited rescue activity, one of two ryanodine receptor antagonists tested (JTV-519, but not Dantrolene) suppressed the AP’s elevated ER stress response (**Figure 6D**).

**Figure 6.**
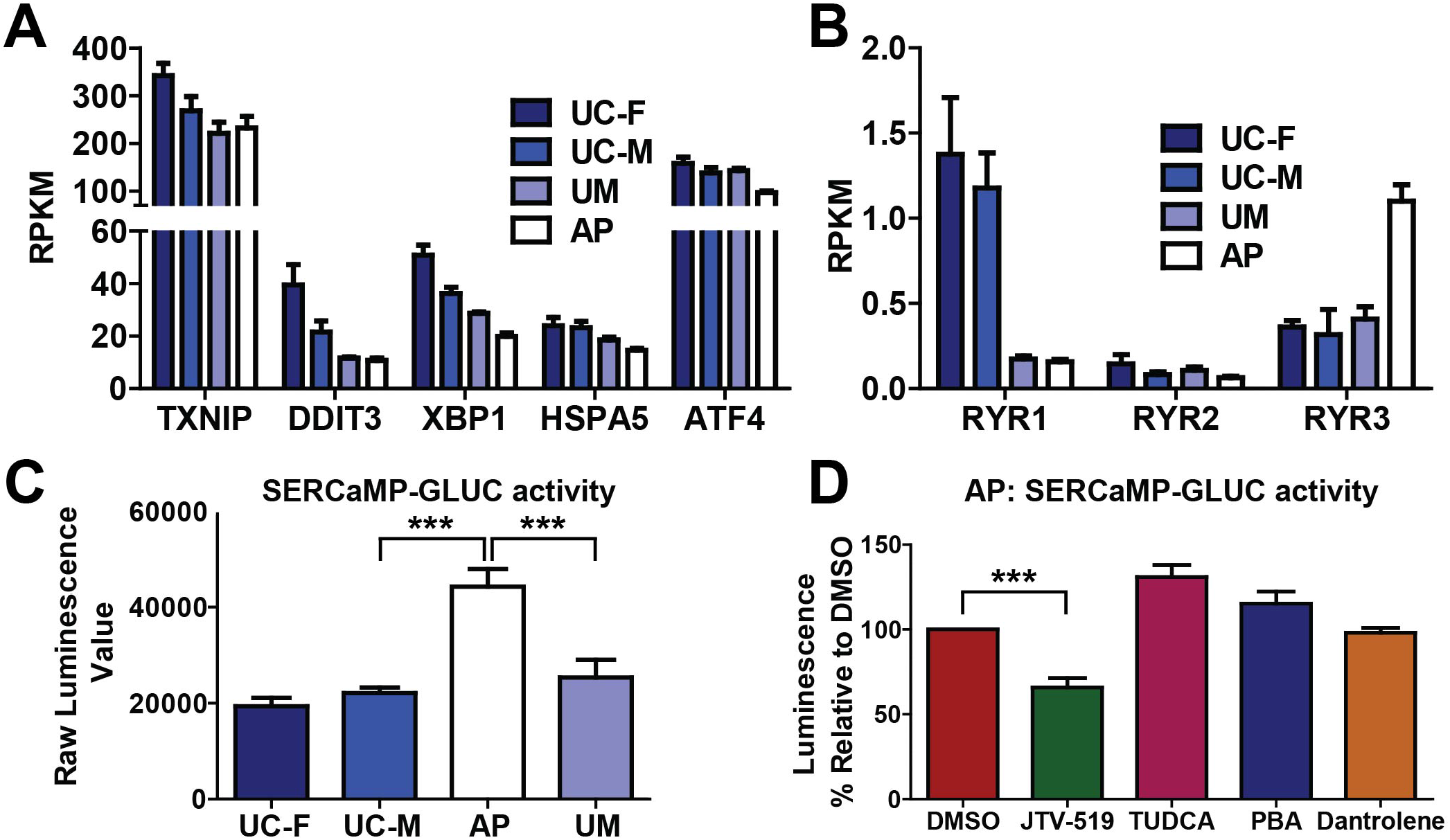
AP-derived cINPCs exhibit elevated ER stress, which is rescued by the ryanodine receptor antagonist JTV-519. (A-B) Analysis of differential gene expression between the models did not identify a significant difference in expression of ER chaperones and ER stress genes in (A), while (B) the AP-derived cINPCs exhibited increased expression of the brain-specific ryanodine receptor RYR3. RNA-seq data is derived from four biological replicate experiments (n=4), which used one clonal line per model as described in the methods. (C) Expression of a Secreted ER Calcium Monitoring Protein Gaussia Luciferase stress sensor (SERCaMP-GLUC) in cINPCs demonstrated elevated ER stress in AP-derived cINPCs. (D) Use of the SERCaMP-GLUC reporter assay in the AP model demonstrated that this elevated ER stress phenotype could be suppressed by treatment with the ryanodine receptor antagonist JTV-519, but not with the chemical chaperones PBA and TUDCA or with another ryanodine receptor antagonist, dantrolene. Four independent biological replicate experiments (n=4) were performed using two clonal lines for the AP and UM models and one clonal line for the UC models. p-values \*\*\**P* < 0.001 were determined by an unpaired t-test.

### Electrophysiological characterization of cortical neurons derived from the AP, UM, and UC-M models

To assess whether neuronal function was altered in these models, cExNPCs and cINPCs were labelled with synapsin-GFP and synapsin–RFP, respectively, these NPCs were differentiated as neuroid co-cultures, and further neuronal maturation was obtained by replating these neuroids on a rat cortical astrocyte feeder layer, as described in the methods and shown in **Supplemental Figure S7A**. Whole-cell voltage and current clamp recordings were then obtained separately from excitatory and inhibitory neurons differentiated from the UC, UM, and AP lines (18 to 27 cells assessed per group). A summary of the electrophysiological analyses performed is in **Supplemental Table S6**. Visual inspection of the cultures suggested that the UM derived neurons were consistently larger than the UC or AP cells, and this was confirmed by cell soma diameter measurements (**Figure 7A; Supplemental Figure S7B**). This was associated with higher UM cell capacitance (proportional to surface area) and lower input resistance (**Figure 7B**). Voltage steps from −80 to 0 mV evoked a fast peak of inward current mediated by tetrodotoxin-sensitive sodium channels, followed by steady-state outward current mediated by voltage-gated potassium channels. Comparison among the three lines revealed that both the UM and AP had a lower outward current density than the UC (**Figure 7C**). In addition, 2-way ANOVA analysis among all groups indicated that, relative to cExNs, cINs had a generally higher input resistance and more depolarized membrane potential for peak inward current (**Figure 7D**). Interestingly, although separate recordings were generated and analyzed for cExNs and cINs across the three sample types, most significant differences were commonly observed in both cEX and cINs. Therefore, we combined the cIN and cExN recording data points for the results reported in **Figures 7–8**. A complete summary of the physiological recordings performed is also provided in **Supplemental Table S6**.

**Figure 7.**
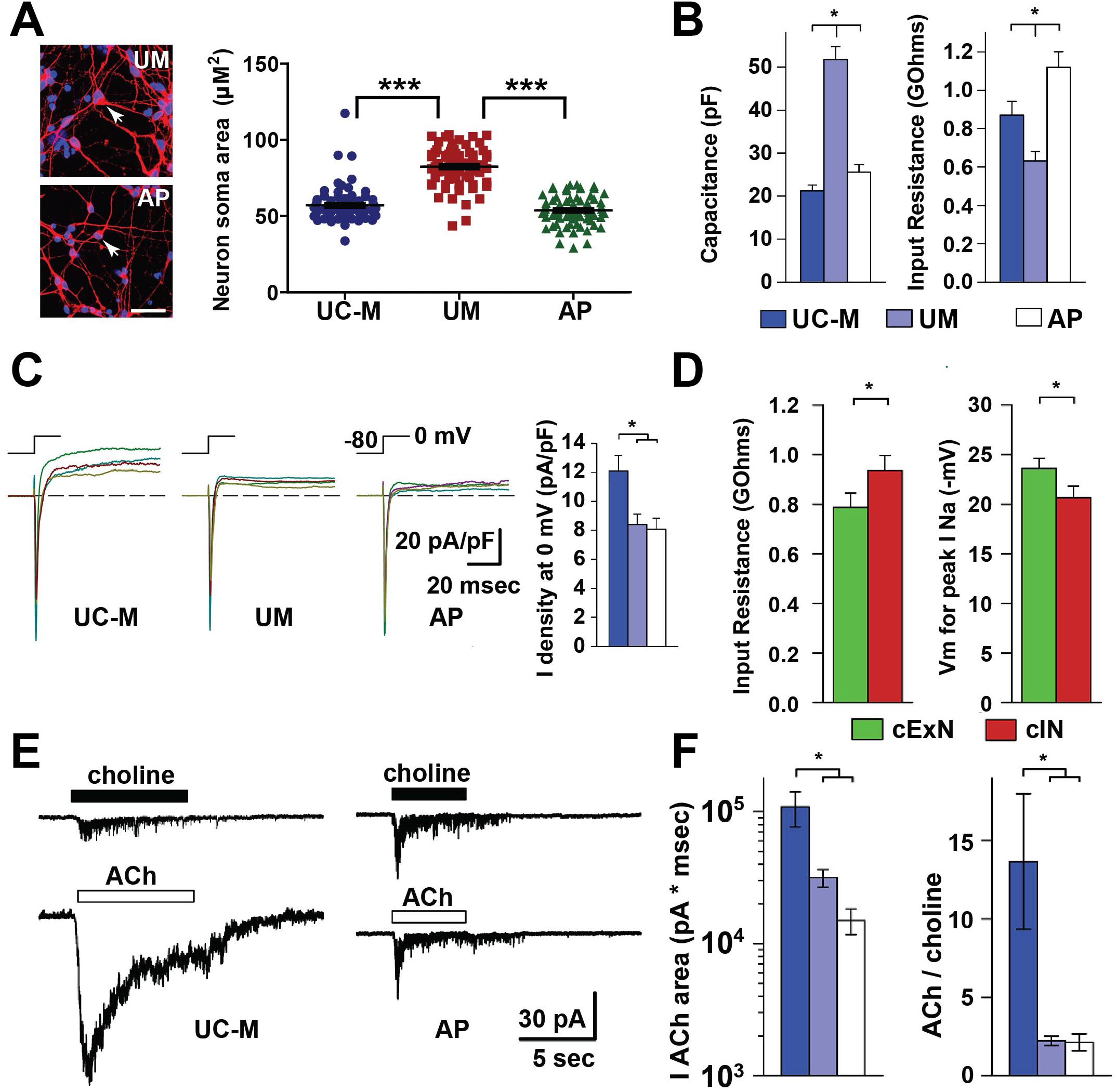
Functional differences in neurons derived from these models were assessed by voltage clamp analysis. (A) Quantitation of neuron soma area revealed a larger soma size in the UM-derived neurons. Left: representative confocal images comparing the AP and UM models, with arrows highlighting a cell soma; Right: quantitation of neuron soma area for the three models. Data is derived from three biological replicate experiments (n=3) using two clonal lines for the AP and UM models, and one clonal line for the UC-M model. (B) UM-derived neurons exhibit significantly higher cell capacitance and lower input resistance than UC-M- or AP-derived neurons. (C) Steady-state outward current density at 0mV was significantly greater for UC-M-than for UM- or AP-derived neurons. Whole-cell inward and outward current density for currents were evoked by a voltage step from −80 to 0 mV. (D) Collectively, iPSC-derived cINs from all three subject-derived models exhibited higher input resistance and less negative voltage for peak inward current (I Na) than cExNs. (E) Whole-cell currents were evoked by 500 μM Choline or ACh in neurons, with representative data shown for the UC-M (left) versus AP (right) models. (F) The integrated area of ACh-evoked current and the ACh/choline ratio for integrated current were both significantly larger for UC-M-derived neurons, by comparison with UM- and AP-derived neurons. Data were derived from three biological replicate experiments (n=3), which used two clonal lines for the AP and UM models, and one clonal line for the UC-M model. p < 0.05, was determined by 2-way ANOVA with post hoc Student-Newman-Keuls test. A complete summary of the physiological recordings performed is also provided in **Supplemental Table S6**.

**Figure 8.**
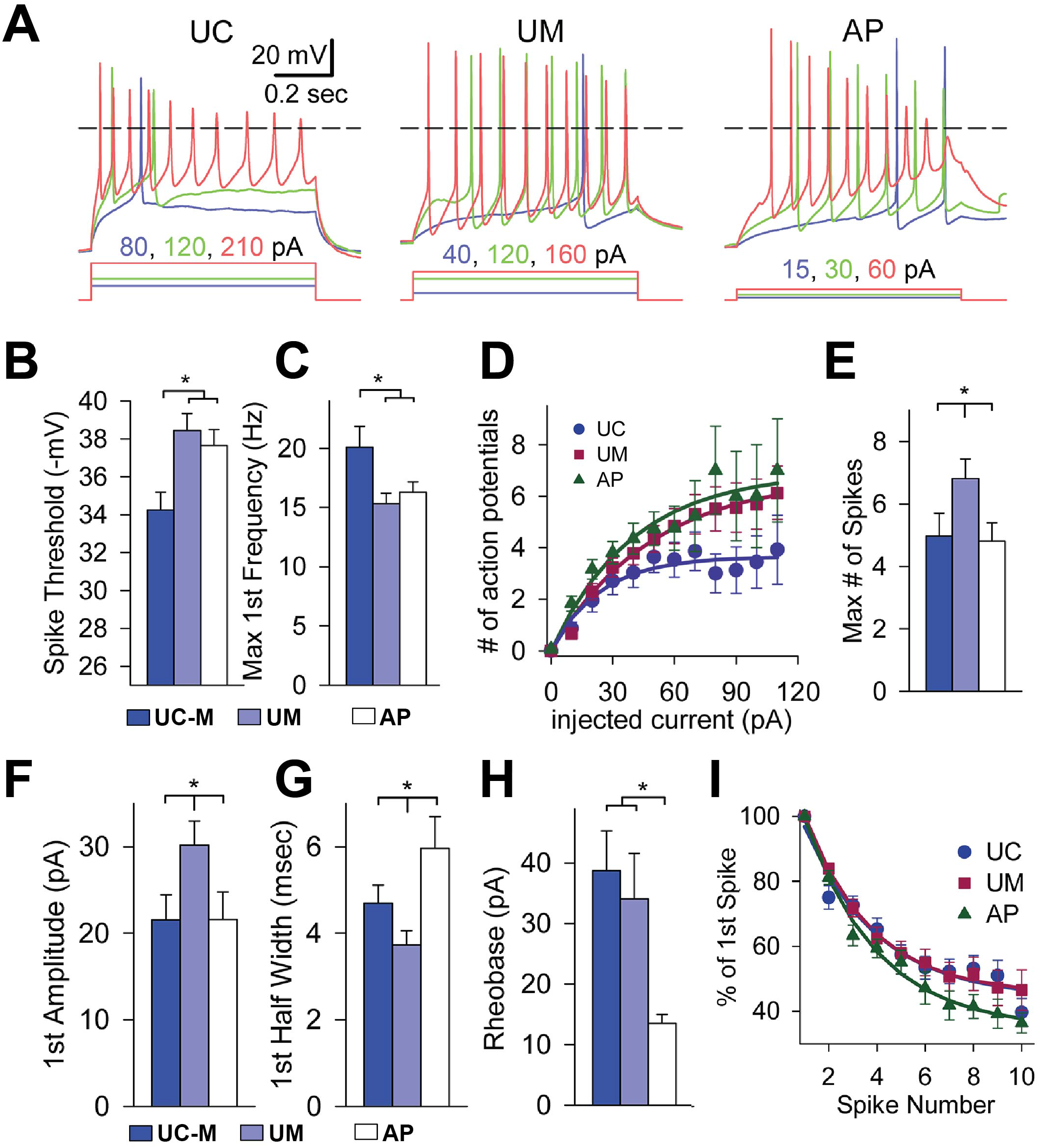
Functional differences in neurons derived from these models were assessed by current clamp analysis. (A-C) Action potentials elicited by three different 800 msec depolarizing current injections in iPSC-derived neurons from the UC, UM, and AP models. Recordings under current clamp revealed (B) a more depolarized threshold for action potential initiation and (C) a higher initial spike frequency in UC-M-derived neurons, as compared to both the UM- and AP-derived models. (D) Exponential fits (smooth curves) indicate significantly fewer action potentials elicited by current injections up to 120 pA in neurons derived from the UC-M model, by comparison with the UM and AP models (p< 0.05, F-statistic). (E-G) UM-derived neurons exhibited a significantly higher (E) maximal number of spikes and (F) first spike amplitude, and a (G) briefer first spike halfwidth than UC or AP neurons. (H-I) Compared to UC- and UM-derived neurons, AP-derived neurons exhibited (H) a substantially lower rheobase and (I) a more significant decline in action potential peak amplitude with each succeeding spike (A and I). p < 0.05 was defined by 2-way ANOVA with post hoc Student-Newman-Keuls test. A complete summary of the physiological recordings performed is also provided in Supplemental Table S6.

Because *CHRNA7* encodes the α-7 nicotinic acetylcholine (ACh) receptor subunit (16), we measured the integrated current elicited by brief exposure to 500 μM ACh. Most cells were also tested with 500 μM choline, which is a relatively selective agonist for homomeric α7 receptors (39), with much weaker activity at heteromeric α7β2 receptors, or other nicotinic receptors expressed in the central nervous system (40, 41). Interestingly, the UC neurons exhibited a significantly larger response to ACh compared to both the UM and AP, while the ACh to choline ratio was also significantly larger for UC neurons, versus both the UM and AP (**Figure 7E-F**, note the log scale in F). These data suggest that homomeric α7 receptors may mediate a higher proportion of the total ACh-evoked current in neurons derived from both the UM and AP, by comparison with the UC-derived neurons. The UM and AP also exhibited other shared alterations of neuronal function. These included a less depolarized threshold for action potential initiation and a lower initial spike frequency than the UC cells (**Figure 8A-C**), while the UM and AP cells also fired a larger number of action potentials than the UC cells for current injections up to 120 pA (**Figure 8D**). In addition to these shared functional alterations, the UM and AP neurons also exhibited some unique functional differences: UM neurons fired a larger maximal number of spikes during an 800 msec depolarizing pulse than neurons derived from UC or AP cells (**Figure 8E**) and also exhibited a larger peak amplitude and shorter half width to the first spike than either UC or AP cells (**Figure 8F-G**), while AP-derived neurons exhibited a substantially lower rheobase (the minimal depolarizing current required to reach threshold; **Figure 8H**) and also exhibited a more significant decline in action potential peak amplitude with each succeeding spike (**Figure 8A, 8I**), compared with both the UM and UC. Therefore, by contrast with the AP-specific cellular and molecular phenotypes described above, the AP and UM shared a number of functional anomalies not seen in the UC model, while also exhibiting some unique characteristics.

## Discussion

Like many CNVs involving microduplication, duplication of genes in the 15q13.3 interval causes a range of psychiatric phenotypes, with this patient population exhibiting widely varied levels of clinical phenotypic penetrance. The extent to which duplication of the *CHRNA7* gene alone, versus other genes in this interval, contributes to these phenotypes, and the basis of its variable penetrance remain largely uncharacterized. Therefore, here we derived iPSC models from individuals in a pedigree where multiple individuals carry the same 15q13.3 CNV involving only duplication of the *CHRNA7* gene, but exhibit substantial differences in affectation. We compared clonal iPSC models derived from a mother with no clinical IDD phenotypes (the UM), her son, who has multiple IDD clinical phenotypes (the AP), and unrelated, unaffected male and female controls (UC-M/UC-F) lacking *CHRNA7* duplication, deriving cortical excitatory (cExN) and inhibitory (cIN) neurons from these iPSC lines to model how neuronal development and function are affected by *CHRNA7* duplication. This approach enabled us to compare potential contributors to differential phenotypic penetrance of *CHRNA7* duplication in the context of a partially shared genetic background. We found that, while both the AP and UM had elevated CHRNA7 expression levels relative to controls, the AP model exhibited a range of neurodevelopmental phenotypes not seen in either the UM or UC-M/F models. These included reduced neurite extension and length, diminished neuronal migration and formation of VGAT-expressing synaptic punctae in cINs, elevated ER stress, and a corresponding reduction of gene expression in related molecular pathways and processes. Intriguingly, many of the same pathways with reduced expression in the AP instead exhibited increased expression in the UM model, by comparison with both the AP and controls. This compensatory molecular signaling could have overcome some consequences of the genetic liability of *CHRNA7* duplication, and may have enabled the UM model to overcome the neurodevelopmental impairment likely to be related to the AP’s clinical phenotypic penetrance. However, examination of electrophysiological function in AP- and UM-derived neurons revealed shared functional anomalies, not seen in controls, including increased action potential firing and enhanced choline responsiveness. These functional changes are consistent with elevated CHRNA7 channel activity and altered ligand sensitivity and indicate some common alterations of neuronal function conferred by *CHRNA7* duplication in the AP and UM models that, if also manifested *in vivo*, are nonetheless insufficient to trigger clinical phenotypes in the UM.

This study generally illustrates the potential for patient-derived neuronal models to identify cellular and molecular correlates and potential contributors to differing levels of clinical phenotypic penetrance. Although many other CNVs involving duplication of genomic intervals likewise exhibit highly variable phenotypic penetrance, the basis of such differential penetrance is not well understood. Here, we could distinguish both contributors that tracked with differential levels of affectation, and those common to multiple carriers of a genetic liability, regardless of clinical phenotype. We could also identify compensatory changes in molecular signaling that may have contributed to a lack of clinical phenotype in the UM. These findings indicate that patient-derived iPSC models may have utility for linking genetic liabilities and differing levels of phenotypic penetrance stemming from those liabilities to particular cellular, molecular, or functional alterations that may be prior or ongoing contributors to the clinical phenotype. These phenotypes then provide a basis and experimental platform for conducting screens to define molecular and pharmacological agents that can reverse affectation-linked phenotypes, as we did in identifying distinct pharmacological agents that could rescue both ER stress and impaired cIN migration in the AP model.

Congruent with the findings of Gillentine et al. (31), we observed elevated ER stress in iPSC-derived NPCs carrying *CHRNA7* duplication in the AP but not the UM model. However, the ER stress phenotypes observed in these two studies differ substantially. Trafficking of nicotinic acetylcholine receptors in the cell involves assembly of five subunits into a pentameric receptor in the endoplasmic reticulum (ER), followed by receptor trafficking to the plasma membrane (42, 43). In Gillentine et al., elevated ER stress was attributed to the unfolded protein response (UPR), based upon increased expression of several ER chaperone- and UPR-related marker genes (31). By contrast, our gene expression profiling by RNA-seq did not reveal altered expression of these or other UPR-related markers in the AP, relative to controls. However, altered calcium homeostasis is another aspect of ER stress (44): to maintain calcium homeostasis, calcium is released from the ER via several mechanisms, one of which involves activation of ryanodine receptor signaling in response to changes in cytoplasmic calcium levels through a calcium-induced calcium release (CICR) mechanism (45). Our gene expression profiling revealed elevated levels of both the calcium receptors CACNA1A, CACNA1B, and CACNA2D2, and of the ryanodine receptor RYR3, which has brain-enriched expression (45), suggesting potentially altered calcium homeostasis. Accordingly, a reporter assay detected elevated ER stress-linked CICR in the AP, relative to both the UM and both unrelated (UC) controls, and we found that the ryanodine receptor antagonist JTV-519 selectively reduced the AP’s ER stress response to baseline conditions, while antagonists of the UPR did not. These results further implicate altered calcium homeostasis rather than UPR pathway activation as the major contributor to the AP’s elevated ER stress in our work here. Another major difference between findings in these studies is that the NPCs in Gillentine et al. exhibited characteristics consistent with reduced CHRNA7 protein trafficking to the cell membrane and reduced CHRNA7 channel activity. By contrast, when we differentiated NPCs into neurons in our study and conducted electrophysiology, we instead found that both *CHRNA7* duplication carriers (the AP and UM) displayed hallmarks of increased CHRNA7 channel activity, including an increased rate of action potential firing and altered ligand sensitivity. Therefore, although both studies observed elevated ER stress in models with *CHRNA7* duplication, we used reporter assays and pharmacological rescue to demonstrate that, for the AP model, this involved altered calcium homeostasis rather than the UPR, and that the consequence of *CHRNA7* duplication in the AP and UM was increased, rather than diminished channel activity. It will be interesting to assess these phenotypes in additional models of *CHRNA7* duplication in future work, to determine whether the effects of *CHRNA7* gene duplication on channel function indeed vary between models derived from different subjects and could be related to other aspects of a particular model, such as interactions with the patient’s genetic background, specific clinical phenotypes, or degree of phenotypic penetrance.

Functional assessment of neurons differentiated from our models by electrophysiology revealed a number of electrophysiological abnormalities common to both individuals with *CHRNA7* duplication (the AP and UM), regardless of the level of affectation. The CHRNA7 receptor can form either a homomeric channel consisting of five α7 subunits, or a heteromeric form comprised of both α7 and β2 subunits. These channels exhibit differential responsiveness, with homomeric α7 receptors having selective sensitivity to choline, while heteromeric receptors are more responsive to acetylcholine (39). Here, both the AP- and UM-derived neurons exhibited elevated CHRNA7 receptor responsiveness to choline and diminished receptor responsiveness to acetylcholine, relative to controls. This is congruent with the possibility that *CHRNA7* duplication, which increased CHRNA7 expression levels in the AP and UM models, may have increased the proportion of homomeric α7 channels, altering electrophysiological function and ligand sensitivity. A number of other electrophysiological abnormalities were also shared by UM- and AP-derived neurons, by comparison with UC-derived neurons, including reduction in the outward potassium (K) current and generation of an increased number of action potentials. Together, these results suggest that increased channel activity may enhance entry of calcium, sodium, and potassium ions into neurons in both the AP and UM models, with these elevated ion concentrations potentially increasing action potential formation in both models with 15q13.3 duplication.

In this study, the AP model exhibited a wide range of neurodevelopmental anomalies, including increased NPC proliferation but reduced neurite outgrowth and neurite length and concordant dysregulation of genes mediating axonal guidance, gap junctional connectivity, and neuritogenesis, with most genes in these pathways exhibiting reduced expression in the AP relative to controls. Altered NPC proliferation, neuronal differentiation, neurite extension, neuronal maturation, and synapse formation have also been observed in other iPSC models derived from ASD patients (18–20, 23, 27, 46–49) and/or in clinical studies of ASD and other neuropsychiatric disorders (47, 50–53). These findings are intriguing since, although both the AP and UM share the same *CHRNA7* duplication, only the AP exhibited these neurodevelopmental alterations, which may be contributors to his ASD pathogenesis. Genes dysregulated in the UM were linked to many of the same GO terms as those seen in the AP versus control comparisons, including axonal guidance, integrin and gap junctions, and behavior- and nervous system development. However, many genes in these networks were up-regulated in the UM relative to both the AP and UC controls, suggesting that a cell-intrinsic compensatory effect may have normalized some corresponding neurodevelopmental processes in the UM model, potentially contributing to the lack of clinical phenotypes seen in this subject.

Alterations in GABAergic neurotransmission are implicated in ASD affectation and may result from disruptions of synaptic development, resulting in imbalanced excitatory and inhibitory neuronal activity in the cortex (37, 54). Since most *in vitro* modeling studies characterize neurodevelopmental phenotypes only in excitatory, glutamatergic neurons (18, 27, 55, 56), the extent to which GABAergic inhibitory neurodevelopment is affected in iPSC models of ASD remains largely unexplored. Here, we observed multiple phenotypes only in the AP model’s cINs but not cExNs, suggesting that assessing phenotypes in both excitatory and inhibitory neurons may be beneficial for identifying neuronal cell type-specific alterations of neurodevelopment that may contribute to ASD. For example, we observed reduced formation of VGAT expressing punctae in the AP-derived cINs, by comparison with both the UM and UC-M-derived cIN. This deficit in production of VGAT expressing punctae could contribute to altered cIN function, potentially impairing cIN synaptic activity to disrupt the balance between excitatory and inhibitory neuronal function in the AP. In the cerebral cortex, inhibitory neurons constitute a minority population (~20%) (57, 58), are specified in sub-cortical progenitor territories, and undergo a long-range tangential migration to their targets in the cortex. Migration of these and other neurons is a critical aspect of brain development (59, 60). Accordingly, here, the AP exhibited decreased cIN migration by comparison with controls, as well as diminished expression of corresponding suites of genes involved in regulating neuronal migration. Other studies of patients with both ASD and other neuropsychiatric disorders have likewise revealed evidence of abnormalities in neurogenesis, neuronal migration, and neuronal maturation (2, 61–63). Furthermore, Wnt signaling can contribute to neuronal migration, and we found here that the Wnt agonist CHIR-99021 could partially rescue the AP model’s migration deficit, while molecular analysis likewise revealed reduced expression of many genes involved in Wnt signaling in the AP model. Modulating Wnt signaling has been explored as an option for ASD treatment; however, as both reduced and elevated Wnt activity have been implicated in ASD-related behavioral and cognitive alterations, further studies are required to stratify how Wnt pathway biomarkers are altered in IDDs including ASD (64–66).

## Limitations

This work aimed to use human subject-derived iPSC models to understand how *CHRNA7* duplication affects neuronal development and function and to identify potential contributors to the wide range of variability in phenotypic penetrance among 15q13.3 duplication carriers. As this study design involved comparisons of family members with the same 15q13.3 duplication but with differential affectation, age and sex represent potential confounds for data interpretation. While ASD is more prevalent in males than females in the population, increasing evidence suggests that females often require a higher load of genetic liability to exhibit clinical phenotypes (67). However, in this case the relationship of differential affectation to sex differences is unclear, as 15q13.3 is autosomal rather than sex chromosome-linked, and current modeling approaches for female iPSC lines cannot recapitulate the random X-chromosome inactivation that occurs during development of female somatic tissues including the brain (68, 69). However, in our RNA-seq analysis, fewer than 3% of the DEGs identified were either X chromosome linked or have been shown to exhibit sex-biased expression in human brain (70, 71). Age is another unavoidable confound in studies that involve modeling multiple subjects in a single pedigree. It is currently unclear whether iPSC lines derived from old donors versus young donors exhibit differences in their potential to undergo differentiation or senescence, and this issue is controversial (72, 73). To address these confounding factors, we performed variancePartition analysis on our RNA-seq data. This analysis indicated that differences between samples were predominantly driven by the subject identity and genetic background, while age and sex were minimal contributors to DEG identification. Another confounding variable in many iPSC modeling studies is differences in genetic background. In this study, the AP and UM have a partially shared genetic background, including the same 424 kilobase duplication at chromosomal location 15q13.3. The large size of this duplication precludes CRISPR-mediated correction, which could be useful to define duplication-linked phenotypes on an isogenic background. In addition, we were unable to access biological material from the father, who could have served as an unaffected male first-degree relative control. However, despite these limitations, this work could still identify cellular, molecular, and functional signatures that differed in clinically affected versus unaffected individuals carrying the same IDD contributory genetic duplication on a partially shared genetic background, as well as defining functional neuronal differences that distinguished these models (AP and UM) from unrelated male and female controls that lacked *CHRNA7* duplication.

## Conclusions

It is often challenging to identify variables that contribute to wide differences in clinical phenotype and penetrance among individuals carrying duplications in the same genomic region. These human genetic anomalies often cannot be recapitulated in rodent models, making them less accessible for experimental study. Furthermore, in cross-comparisons between human cellular models, both the particular genomic interval and which gene or genes are involved, as well as heterogeneous genetic background contributors present in unrelated individuals, can confound such cross-comparisons. Here, the use of both comparisons between related individuals with the same duplication but with variable affectation, and with lines derived from unrelated control subjects lacking this genetic anomaly, was informative in defining a number of affectation-linked neurodevelopmental deficits and an ER stress phenotype exhibited specifically by the AP-derived models, some of which may contribute to his severity of affectation. We also identified potentially compensatory up-regulation of some of the same developmental regulatory pathways in the UM model, which could contribute to a lack of neurodevelopmental phenotypes in this model and of clinical phenotypic penetrance in this subject. In addition, and despite their variable affectation, both models carrying *CHRNA7* duplication exhibited overlapping neuronal functional abnormalities. Therefore, these findings highlight the potential for iPSC models to identify cellular and molecular anomalies linked to either the presence versus absence of an IDD-linked CNV, such as 15q13.3 duplication, as well as to differences in phenotypic penetrance among multiple individuals carrying the same CNV.

It would be interesting to examine the extent to which neurodevelopmental deficits, particularly in cINs, occur in and are linked to levels of patient affectation in models derived from other *CHRNA7* duplication carriers, to determine whether some phenotypes identified here frequently predict clinical severity in this patient population. It would also be intriguing to know whether these phenotypes are more broadly generalizable indicators of the level clinical phenotypic penetrance in other IDD-contributory CNVs involving genome microduplication, as many other such CNVs exhibit a similarly wide range in clinical phenotypic penetrance as is seen for 15q13.3. If some of the phenotypes identified here represent general hallmarks of and contributors to the level of clinical phenotypic penetrance in CNVs involving microduplication at either 15q13.3 or other susceptible genomic intervals, this would provide a molecular and cellular basis for molecular or pharmacological screening for interventions capable of phenotypic reversal, while these experimental platforms could have utility for developing interventions for treating affected individuals.

## Abbreviations

ASD: Autism Spectrum Disorder
iPSCs: Induced Pluripotent Stem Cells
CNVs: Copy Number Variations
CHRNA7: α-7 nicotinic acetylcholine receptor subunit
ID: Intellectual Disability
IDD: Intellectual and Developmental Disability
ADHD: Attention Deficit and Hyperactivity Disorder
ER: Endoplasmic Reticulum
GABA: Gamma Aminobutyric Acid
RYR: Ryanodine Receptor
UPR: Unfolded Protein Response
cExNs: Cortical Excitatory Neurons
cINs: Cortical Inhibitory Neurons
NPCs: Neural Progenitor Cells
UM: Unaffected Mother
AP: Affected Proband
AB: Affected Brother
UC-M: Unrelated Control-Male
UC-F: Unrelated ControlFemale
GTAC: Genome Technology Access Center
ICC: Immunocytochemistry
RT-qPCR: reverse transcription and quantitative PCR
EBs: Embryoid Bodies
DEG: Differentially Expressed Gene
IPA: Ingenuity Pathway Analysis
PI: Propidium Iodide
PCA: Principal Component Analysis
GO: Gene Ontology

## Ethics approval and consent to participate

Subjects were consented for biobanking and iPSC line generation by the Washington University Institutional Review Board of the Human Research Protection Office under human studies protocol #201409091 (Dr. John Constantino).

## Consent for publication

Consent to publish data was provided by all subjects.

## Availability of data and materials

The RNA-seq data generated during the current study are available in the Gene Expression Omnibus (GEO) repository as Series GSE143908.

## Author’s contributions

K.M. contributed to the study design, carried out all experimentation, analyzed data, and prepared the manuscript. R.P. contributed to the luciferase experimentation and analyzed data. D.B. interpreted cytogenetics data. P.G. and B.Z. performed RNA-seq data analysis. A.B. contributed to the study design. F.U. contributed to the ER assays. J.E.H. contributed to performing electrophysiological experiments, analysis, interpretation, and manuscript writing. J.N.C. contributed to the study design and manuscript preparation. K.L.K. contributed to the study design, data analysis, and manuscript preparation. All authors read and approved the final manuscript.

## Acknowledgments

We thank the family for providing biomaterials for use in this study. We thank the Genome Engineering and iPSC Center (GEiC) at Washington University School of Medicine (WUSM) for deriving the iPSC lines used in this study. We thank the Alvin J. Siteman Cancer Center at WUSM for the use of the Siteman Flow Cytometry Core, which provided self-service Flow Cytometry Analysis. The Siteman Cancer Center is supported in part by an NCI Cancer Center Support Grant #P30 CA091842. We thank the Genome Technology Access Center in the Department of Genetics at WUSM for providing genomic sequencing services. The Center is partially supported by NCI Cancer Center Support Grant #P30 CA91842 to the Siteman Cancer Center and by ICTS/CTSA Grant# UL1 TR000448 from NIH/NCRR. We also thank the WUSM Cytogenetics & Molecular Pathology specialists for providing karyotyping services.

## Figure and Table Legends

**Table 1.** Differential clinical phenotypes and location of copy number variation in a family with 15q13.3 duplication.

