## Supplemental Figures S1-S7 for "Alterations in neuronal physiology, development, and function associated with a common duplication of chromosome 15 involving *CHRNA7*"

#### Supplemental Fig. S1

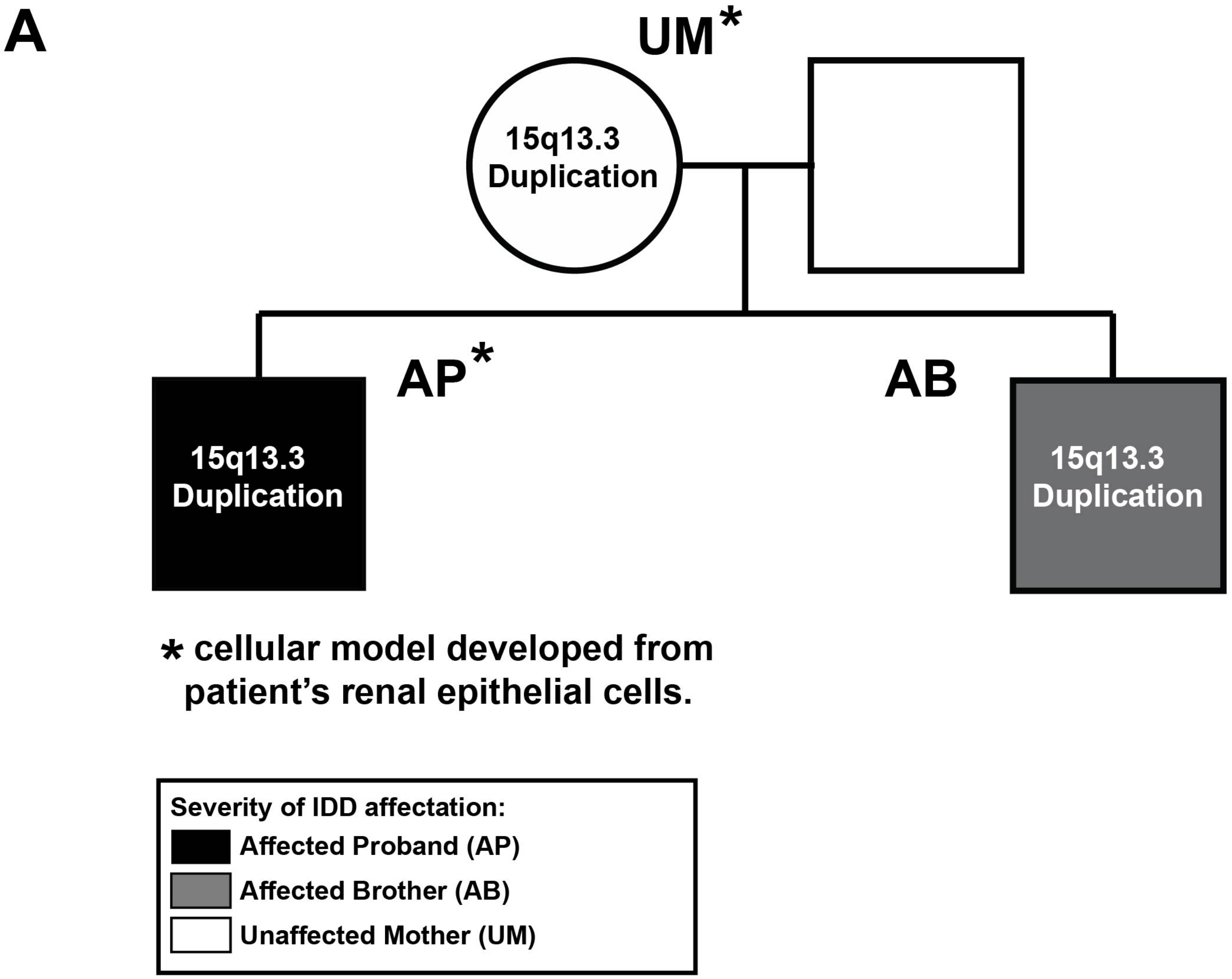


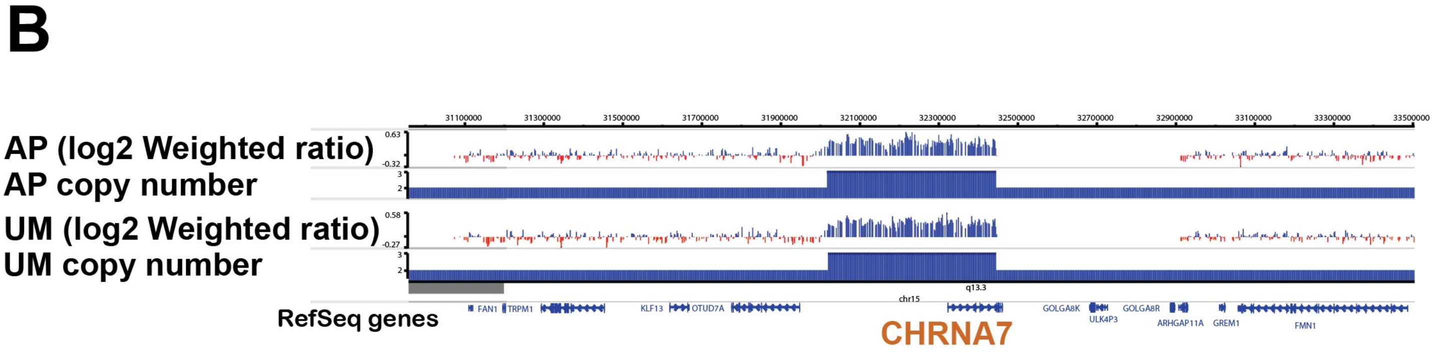

**Supplemental Fig. S2**

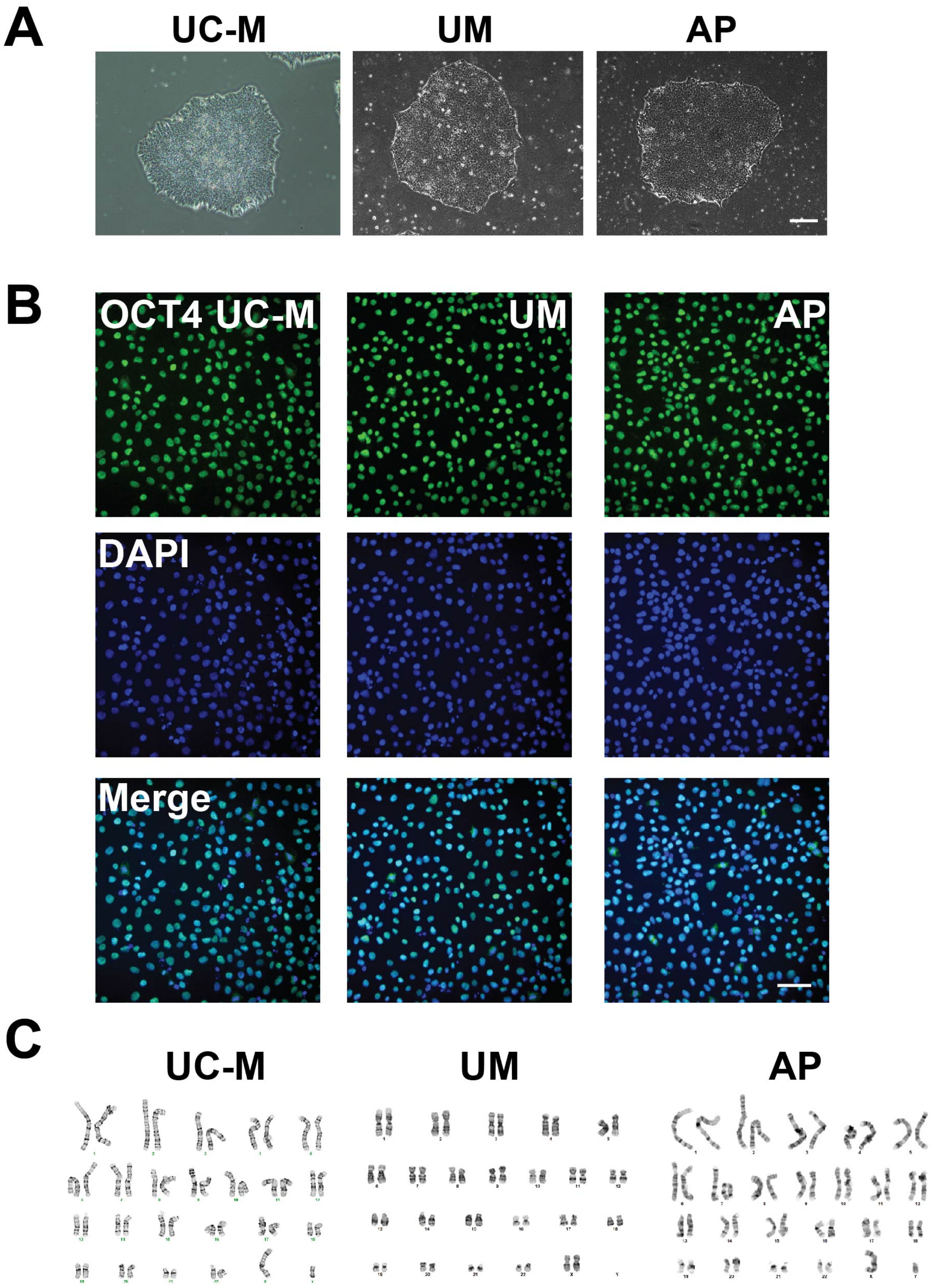

Supplemental Fig. S3

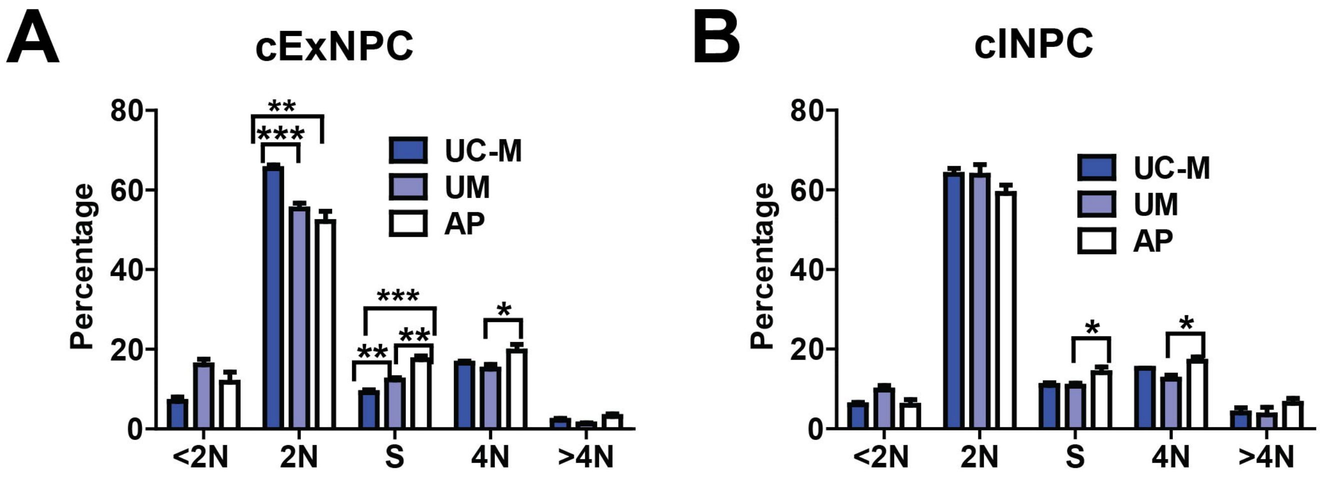

#### Supplemental Fig. S4

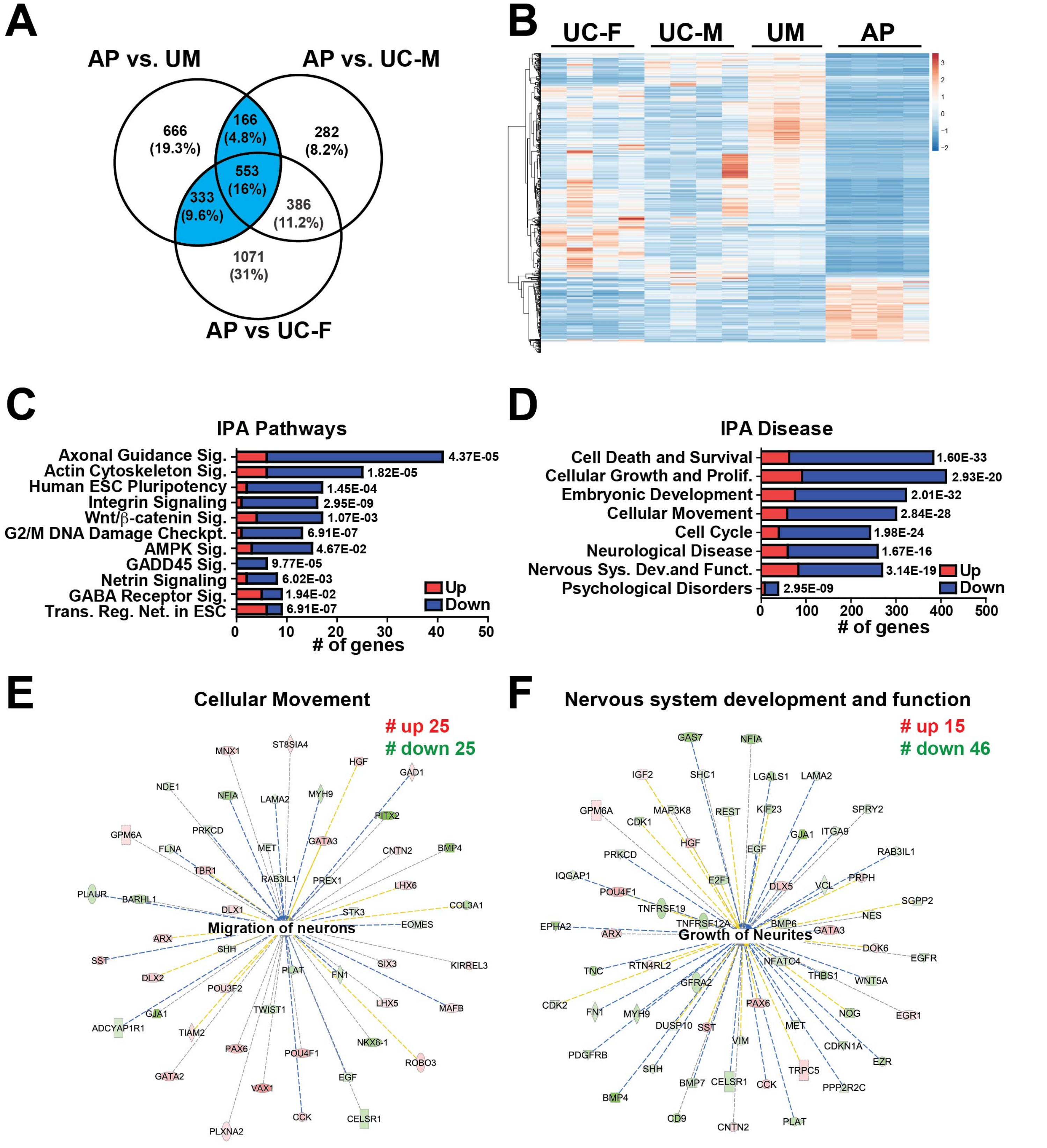

### Supplemental Fig. S5

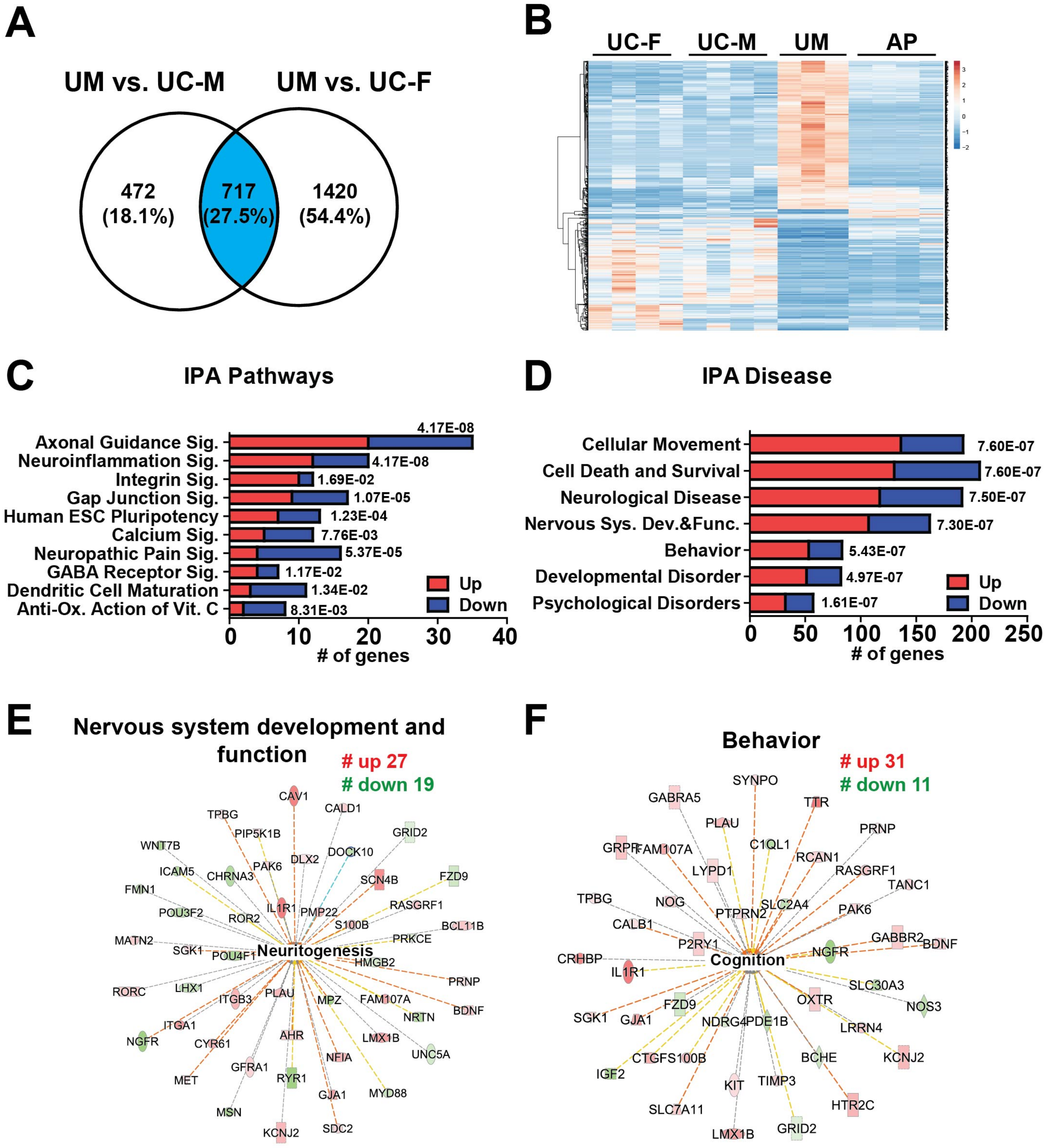

### Supplemental Fig. S6

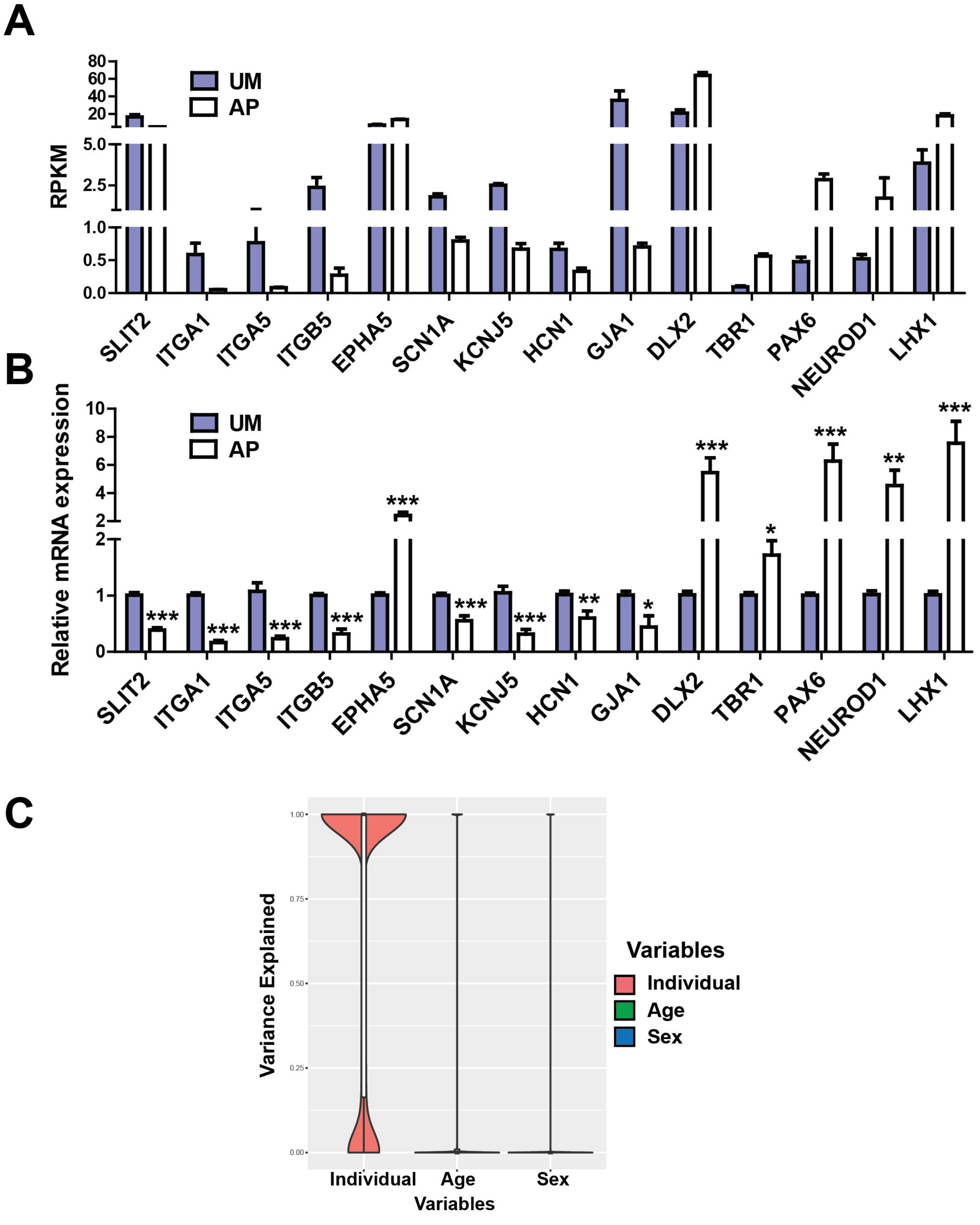

### Supplemental Fig. S7

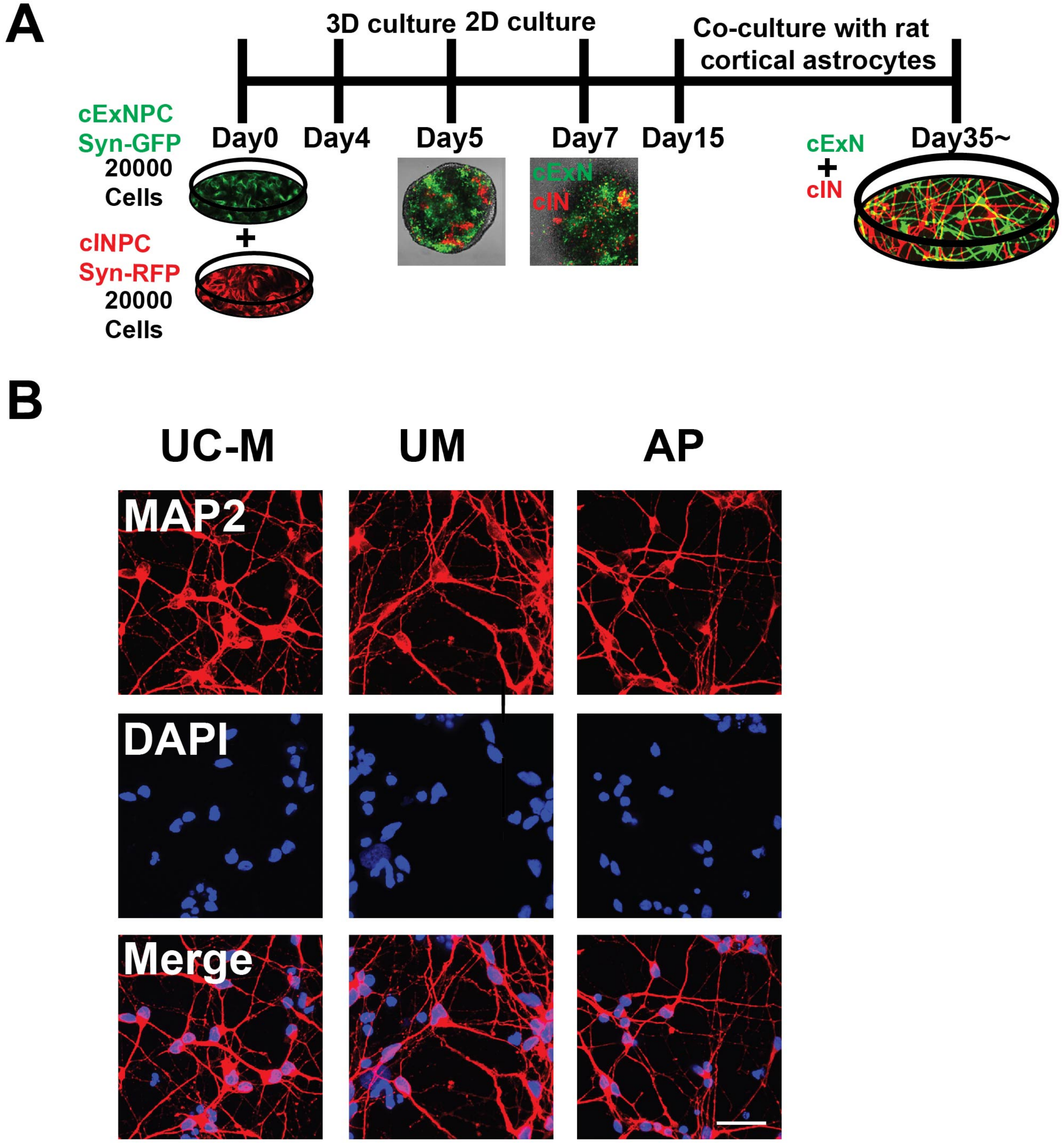
