## Supplemental Figure and Table Legends for "Alterations in neuronal physiology, development, and function associated with a common duplication of chromosome 15 involving *CHRNA7*"

**Supplemental Figure S1. Characteristics of subjects in pedigree modeled here.** (A) The study samples were derived from a pedigree with 15q13.3 duplication, with differential clinical affectation indicated by shading of subjects. The affected proband (AP) is represented in black, his affected brother AB, shown in gray, exhibits subtle autistic traits and has more volatile emotional dysregulation than the AP, while the unaffected mother (UM) and father are shown in white. Renal epithelial cells from the family members indicated (*) were used to derive iPSC models. (B) CNV array data for the AP and UM shows the signal intensity (log2 weighted ratio) and predicted copy number across duplicated region of 15q13.3. The region lacking signal is not covered by the CNV array.

**Supplemental Figure S2. Characterization of iPSC models.** Renal epithelial cell-derived iPSC lines from the UC-M, UM, and AP subjects (A) exhibit normal human stem cell colony morphology in bright field images (scale bar = 250 µm), (B) express the pluripotency marker OCT4/POU5F1 (scale bar = 150 µm), and (C) have a normal karyotype.

**Supplemental Figure S3.** **Cell cycle analysis of neural progenitor cells (NPCs).** (A) cExNPCs and (B) cINPCs were stained with propidium iodide for DNA content and analyzed by FACS. Percentages of cells in each phase of the cell cycle were quantified for each model. Values shown are from seven independent biological replicate experiments (n=7), using two clonal lines for the UM and AP, and one clonal line for UC-M. p-values **P* < 0.05, ***P* < 0.01, ****P* < 0.001 were determined by an unpaired t-test.

**Supplemental Figure S4. Differentially expressed genes in the AP, by comparison with all three other models.** (A) Venn diagram shows numbers of differentially expressed genes (DEGs) obtained from pairwise comparisons of the AP versus (vs) UM, AP vs UC-M, and AP vs UC-F models. AP-specific DEGs, based upon comparisons to at least two of the other datasets, are shaded in blue. These AP-specific DEGs were further analyzed by: (B) Hierarchical clustering analysis, visualizing comparisons with the other three sample types, and by (C-H) Ingenuity Pathway Analysis (IPA), which identified (C) enriched pathways and (D) disease-related GO terms. In C-D, the number of DEGs enriched for each term present is represented on the x-axis, with red and blue colors indicating up- and down-regulated genes, respectively. *P*-values for each enriched GO term are indicated. (E-F) IPA disease terms enriched in these AP-specific DEGs include gene networks associated with (E) Cellular movement and (F) Nervous system development and function. The numbers of up-and down-regulated genes present in the networks are indicated. Within each network, red and green symbols indicate up- and down-regulated genes respectively, while color intensity indicates the relative degree of differential expression.

**Supplemental Figure S5. Differentially expressed genes in the UM, by comparison with the UC-M and UC-F control models.** (A) Venn diagram shows numbers of differentially expressed genes (DEGs) obtained from pairwise comparisons between the UM vs the UC-M or UC-F models. UM-specific DEGs are shown in blue. (B-F) These UM-specific DEGs were further analyzed by (B) Hierarchical clustering analysis, with comparisons to all three other models shown, and (C) using Ingenuity Pathway Analysis (IPA), which identified UM-enriched (C) pathways and (D) disease-related GO terms. In C-D, the number of DEGs enriched for each term present is represented on the x-axis, with red and blue colors indicating up- and down-regulated genes, respectively. *P*-values for each enriched GO term are indicated. (E-F) IPA disease terms enriched in these AP-specific DEGs include gene networks associated with (E) Nervous system development and function and (F) Behavior. The numbers of up-and down-regulated genes present in the networks are indicated. Within each network, red and green symbols indicate up- and down-regulated genes respectively, while color intensity indicates the relative degree of differential expression.

**Supplemental Figure S6. AP-specific differential gene expression defined by RNA-seq analysis of differentiated cortical neuroids was validated by RT-qPCR and variancepartition analysis.** Genes defined as differentially expressed in the AP, by comparison with the UM, by RNA-seq analysis were selected from top AP-enriched gene networks, including axon guidance molecules, integrins, ion channels, and transcription factors, and were validated by RT-qPCR. (A) RPKM values for these DEGs, as obtained using RNA-seq analysis, were (B) compared with relative gene expression in these models as defined by RT-qPCR. (C) variancePartition analysis indicates the percent variance that is attributable to individuals from whom the samples were procured, and the study subjects’ age and sex. *P* values **P* < 0.05, ***P* < 0.01, ****P* < 0.001 were determined by an unpaired t-test. RT-qPCR analysis was performed using samples obtained from three independent biological replicate experiments (n=3) performed using a second set of clonal lines derived from the AP and UM that was different than the AP and UM clonal lines used for the RNA-seq analysis.

**Supplemental Figure S7.** **Electrophysiological characterization of cExNs and cINs.** (A) Schematic of the differentiation approach used to obtain neurons for electrophysiology. cExNPCs and cINPCs, labelled respectively with Synapsin (Syn)-GFP and Syn–RFP, were differentiated in co-culture as cortical neuroids for 15 days, and then further matured by replating on a rat cortical astrocyte feeder layer. (B) MAP2 staining of dissociated cortical neuroids demonstrated that the UM-derived neurons had increased soma size, as quantified in **Fig. 7A.** (Scale bar = 75 µM).

**Supplemental Table Legends**

**Supplemental Table 1**. (A) Antibodies used for immunocytochemistry, with the dilutions, suppliers, and host species indicated. (B) Number of Ki67-expressing NPCs and total number of DAPI-stained nuclei quantified to define the Ki67-expressing fraction in **Fig. 2F**. For **Fig. 3D**, total number of neurons used for quantitation of neurite length, based upon number of DAPI-expressing nuclei. (C) Indication of which clonal line was used for each biological replicate experiment and the number of biological replicates that were performed to procure the data for each figure panel.

**Supplemental Table 2**. Differentially expressed genes were obtained by pairwise comparisons of normalized RNA-seq expression data from the four models, including calculation of the log2 fold change, FDR corrected p-values (padj), and average RPKM values across the sample types analyzed. See Methods for further information.

**Supplemental Table 3**. Ingenuity Pathways Analysis (IPA) of DEGs specific to the AP, by comparison with the UM. Significantly enriched: (A) Pathway and (B) Disease-related GO terms are shown, with the p-value and the DEGs from which each term was derived.

**Supplemental Table 4**. Ingenuity Pathways Analysis (IPA) of DEGs specific to the AP, by comparison with two or more of the other (UM, UC-M, and/or UC-F) models. Significantly enriched: (A) Pathway and (B) Disease-related GO terms are shown, with the p-value and the DEGs from which each term was derived.

**Supplemental Table 5**. Ingenuity Pathways Analysis (IPA) of DEGs specific to the UM, by comparison with the UC-M and UC-F models. Significantly enriched: (A) Pathway and (B) Disease-related GO terms are shown, with the p-value and the DEGs from which each term was derived.

**Supplemental Table 6.** Comparison of electrophysiological properties of neurons derived *in vitro* from the UC-M, AP, and UM iPSC models is shown for (A) all cells, (B) cExNs, and (C) cINs. (A) Data columns for cells of each genotype show the mean, SEM, and count (# of cells) for each parameter for all cells recorded, including cells from both excitatory and inhibitory induction protocols. Columns to the right show combined data for all of the cells of all three genotypes from excitatory inductions and all of the cells from inhibitory inductions. Light red shading denotes Significant Difference by 2-WAY ANOVA (p<0.05) with genotype (UC-M : AP : UM) and induction protocol (cExN : cIN) as the two factors. (B-C) Values for cExN and cIN cells of each genotype are shown below the table for all cells. Tan shading denotes Significant Difference by 1-WAY ANOVA (p<0.05) with darker tan indicating that one of the genotypes was different from the other two and lighter tan indicating that two of the three genotypes were different from each other. Capacitance and input resistance were determined from 10 mV voltage steps from -80 mV. First spike threshold, amplitude and half-width were determined for the first action potential recorded at threshold. Maximum first frequency and maximum average firing frequency were determined for 800 msec depolarizing steps that elicited the maximum number of spikes under current clamp. Peak inward sodium current, steady-state outward potassium currents and currents evoked by ACh and choline were recorded under whole-cell voltage clamp.
